# Predicting the Immunogenicity of T cell epitopes: From HIV to SARS-CoV-2

**DOI:** 10.1101/2020.05.14.095885

**Authors:** Ang Gao, Zhilin Chen, Florencia Pereyra Segal, Mary Carrington, Hendrik Streeck, Arup K. Chakraborty, Boris Julg

## Abstract

We describe a physics-based learning model for predicting the immunogenicity of Cytotoxic T Lymphocyte (CTL) epitopes derived from diverse pathogens, given a Human Leukocyte Antigen (HLA) genotype. The model was trained and tested on experimental data on the relative immunodominance of CTL epitopes in Human Immunodeficiency Virus infection. The method is more accurate than publicly available models. Our model predicts that only a fraction of SARS-CoV-2 epitopes that have been predicted to bind to HLA molecules is immunogenic. The immunogenic CTL epitopes across all SARS-CoV-2 proteins are predicted to provide broad population coverage, but the immunogenic epitopes in the SARS-CoV-2 spike protein alone are unlikely to do so. Our model predicts that several immunogenic SARS-CoV-2 CTL epitopes are identical to those contained in low-pathogenicity coronaviruses circulating in the population. Thus, we suggest that some level of CTL immunity against COVID-19 may be present in some individuals prior to SARS-CoV-2 infection.

## Introduction

Pandemics caused by infectious pathogens (e.g., viruses, bacteria) have plagued humanity since antiquity. The latest example is the Coronavirus Disease 2019 (COVID-19) caused by the SARS-CoV-2 virus, which has already exacted a heavy global toll on human health and the economy. One key tool in fighting infectious diseases are vaccines. Vaccination has led to the eradication of smallpox from the planet, the near-eradication of polio, and contributed greatly to the reduction in childhood mortality. Indeed, vaccination has saved more lives than any other medical procedure. Bringing the COVID-19 pandemic under control will likely require an effective vaccine. Thus, several urgent efforts to develop vaccines that may protect against infection by SARS-CoV-2 have been launched (Akst, 2020). For example, *Moderna* has announced clinical trials of a messenger RNA-based vaccine that codes for the spike protein of SARS-CoV-2 (Dunn, 2020). Other ongoing efforts involve the use of non-replicating Adenovirus vectors containing the gene that encodes the spike protein of SARS-CoV-2 (Mak, 2020). Both these approaches aim to elicit protective antibody responses. Whether these, and other vaccines in development will elicit protective immune responses, and how durable the protection will be, are not known.

SARS-CoV-2 is a coronavirus of the same family as the viruses that caused Severe Acute Respiratory Syndrome (SARS) in 2003 and Middle East Respiratory Syndrome (MERS) in 2012. Phylogenetic analyses based on available sequences of SARS-CoV-2 suggest that the new virus is most similar to SARS-CoV (Lu *et al*., 2020; Zhou *et al*., 2020). A recent report shows that the nucleocapsid (N), membrane (M), and envelope (E) proteins of SARS-CoV-2 are over 90 % conserved compared to SARS-CoV, and the spike (S) protein is 76 % similar (Ahmed, Quadeer and McKay, 2020). Most effective prophylactic vaccines elicit a potent antibody response directed against the spike proteins of viruses. But, a number of studies show that the antibody response elicited in patients infected with SARS-CoV was protective but relatively short-lived (Liu *et al*., 2006; Mo *et al*., 2006; Tang *et al*., 2011), while T cell responses were durable (Fan *et al*., 2009; Tang *et al*., 2011; Channappanavar *et al*., 2014). For example, Fan et al (Fan *et al*., 2009) showed that most patients who recovered from SARS-CoV have memory T cell responses directed against the virus 4 years after recovery. Tang et al (Tang *et al*., 2011) showed that 6 years after recovery SARS-CoV patients did not have significant amounts of virus-specific circulating antibodies, but had significant memory T cell responses compared to healthy controls. Furthermore, a critical role for virus-specific memory T-cells in broad and long-term protection against SARS-CoV infection has been elucidated in animal models (Zhao, Zhao and Perlman, 2010; Channappanavar *et al*., 2014). Therefore, given the similarities between SARS-CoV-2 and SARS-CoV, it is worth exploring the development of vaccines that may elicit protective T cell responses.

T cells target pathogenic peptides (epitopes) bound to Human Leukocyte Antigen (HLA) molecules. There is a great diversity of HLA genes in the human population, with each individual possessing 6-12 types of alleles (Bui *et al*., 2006). Multiple recent studies have been focusing on discovering potential epitopes of SARS-CoV-2 that can elicit T cell responses. Ahmed et al (Ahmed, Quadeer and McKay, 2020) and Grifoni et al (Grifoni *et al*., 2020) have tried to identify peptides of SARS-CoV-2 that have high sequence identity with SARS-CoV epitopes. However, only a small number of SARS-CoV peptides that are experimentally known to elicit T cell responses in humans are shared by SARS-CoV-2. Moreover, these shared peptides are associated with a limited set of HLA molecules, thus providing poor coverage of the global population. Ahmed et al (Ahmed, Quadeer and McKay, 2020) and Prachar et al (Prachar *et al*., 2020), among other groups (Campbell *et al*., 2020; Nerli and Sgourakis, 2020; Prachar *et al*., 2020), also identified SARS-CoV-2 peptides that are capable of binding to HLA molecules, either based on MHC binding assay results or bioinformatic methods. By doing this, they identified a large pool of SARS-CoV-2 peptides that are associated with diverse HLA molecules, which cover a broad cross-section of the global population. But, binding to HLA molecules does not imply that the peptide epitope will elicit an immunogenic T cell response in humans (Yewdell, 2006).

A number of efforts have aimed to develop bioinformatic tools to characterize the sequences of TCRs in human repertoires, and to follow how particular clones evolve in response to viral infections (e.g. Murugan *et al*., 2012; Minervina *et al*., 2020). By analyzing sequences of TCRs that bind to a panel of epitopes, Glanville et al and Dash et al (Dash *et al*., 2017; Glanville *et al*., 2017) discovered that TCRs that bind to the same epitope often share conserved sequence motifs. They then constructed a sequence-similarity-based clustering algorithm that, with high probability, clusters TCRs with the same epitope specificity together. Our goal here was different.

We aimed to determine the relative immunogenicity of SARS-CoV-2 T cell epitopes in people with diverse HLA alleles by developing a physics-based learning algorithm. To train and test our method we first studied another viral infection, Human Immune Deficiency Virus (HIV), for which well defined data on the relative immunodominance of T cell epitopes is available. The results suggest that our method is more accurate than the immunogenicity prediction tool publicly available on IEDB (Calis *et al*., 2013; Vita *et al*., 2019). Furthermore, the nature of our model suggests that the algorithm should be able to predict relative immunogenicity of peptide epitopes derived from a variety of pathogens.

Therefore, we applied our algorithm to identify immunogenic SARS-CoV-2 peptide epitopes. Our results show that only a fraction of peptide epitopes that are known to bind different HLA molecules is likely to be immunogenic. However, the set of immunogenic peptides still provides broad coverage of the global population. Given the low mutability of the SARS-CoV-2 virus so far, these results suggest that a whole proteome immunogen may be able to elicit potent T cell responses in diverse individuals. We also predict that the immunogenic CTL epitopes contained in the spike protein of SARS-CoV-2 (immunogen in most current vaccine formulations) is unlikely to provide broad population coverage, since these spike epitopes are associated with limited number of HLA alleles. In addition, several predicted immunogenic peptide epitopes derived from the SARS-CoV-2 proteome are identical to those contained in the four human coronaviruses of low pathogenicity (HCoV) that regularly circulate in the human population. This result leads us to suggest that HCoV-specific memory CTL responses that can also target SARS-CoV-2 epitopes in immunogenic fashion may be present in some fraction of the human population even prior to SARS-CoV-2 infection.

Our results for SARS-CoV-2 have not been tested experimentally, but we hope that they will provide a useful guide to prioritize peptides that other investigators may wish to test. We note also that, upon further testing, validation and elaboration, our approach for predicting immunogenicity of T cell epitopes may be useful for future infectious pathogens that will undoubtedly emerge.

## Results

### Model Development and training against clinical data

Li et al (Li *et al*., 2008) carried out a detailed study identifying T cell epitopes targeted by 226 SARS-CoV-infected patients, of which 98 were in an acute state of infection. They found that, both in terms of frequency and magnitude, the CTL response was dominant compared to CD4 T cell responses. Therefore, here we will focus on CTLs and the relative immunogenicity of peptide epitopes that they may target in humans.

CTLs bind to short peptides, about 9 – 11 amino acids in length, displayed in complex with HLA class I molecules. Such short motifs do not contain any long-range information about the genome of the organism from which they are derived (Kosmrlj *et al*., 2008; Butler, Kardar and Chakraborty, 2013). Therefore, the ability to predict relative immunogenicity of peptide epitopes derived from the genome of one virus in persons with a given HLA allele is likely to allow prediction of immunogenicity of epitopes derived from another virus’ proteome.

Our model for immunogenicity of CTL peptide epitopes is inspired by studies aimed to predict immunogenicity of cancer neo-antigens for immunotherapy (Luksza *et al*., 2017). We wish to predict the peptide immunodominance hierarchy in people with different HLA genes. For that purpose, we define a “CTL response metric” A(s, M), where *s* is the sequence of the peptide whose relative immunogenicity we wish to predict for a person with the HLA allele, *M*. This metric is the product of three terms, and is written as follows:

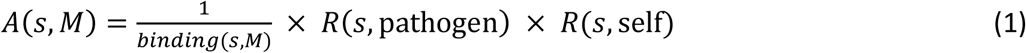

Each of the terms above reflects a different physical phenomenon. The binding term,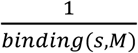 is a measure of the probability that the peptide *s* can be processed, bound to, and displayed by HLA molecule *M*. Machine learning approaches have been trained on many measurements of peptide presentation by different HLA molecules, and a resulting method, netMHCpan4.0 (Vanessa and Nielsen, 2017), can make reasonable estimates of *binding*(*s, M*) for many alleles as the percentile rank of the elution-ligand score. We next posit that whether or not a peptide is targeted by human CTLs should correlate with how similar its sequence is to peptides derived from diverse pathogens that are known to elicit a CTL response in humans (listed in the IEDB database (Vita *et al*., 2019); Methods). The term, *R*(*s*, foreign) in Eq. 1 is the sequence similarity of peptide *s* to these pathogen-derived peptides. We define *R*(*s*, foreign) mathematically as the number of foreign peptides whose alignment score with *s* is larger than a threshold value, *a*_*pathogen*_ :

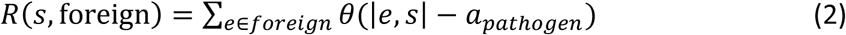

Here, *e* is a pathogenic peptide in the database, |*e, s*| is the alignment score of *e* and *s*, which is determined by the BLOSUM62 based Smith-Waterman alignment method (as used by Luksza et al (Luksza *et al*., 2017)), and *θ* is the step function. A higher alignment score means that the biochemical properties of the two peptides are more similar to each other.

T cells develop in the thymus, where they are exposed to HLA-bound peptides derived from the host’s proteome. For a thymocyte to mature into a peripheral T cell, it must bind to at least one of these peptides with an affinity that exceeds a threshold value, and not bind to any of them with an affinity that exceeds a higher threshold value (Daniels *et al*., 2006). In past studies (Kosmrlj *et al*., 2008, 2009; Košmrlj *et al*., 2010; Butler, Kardar and Chakraborty, 2013), we developed a mechanistic understanding of how thymic development shapes the pathogen reactivity of the T cell repertoire in an organism. Extending these studies leads us to the seemingly counter-intuitive conclusion that T cells that bind to human peptides more strongly will also be likely to bind more strongly to pathogen-derived peptides. One simple way to understand this is by examining data in mice, which show that more self-reactive T cells are statistically enriched in more hydrophobic amino acids at residues that contact the HLA-bound peptides (Stadinski *et al*., 2016). This is because TCR binding to HLA-bound peptides creates an interface from which water must be partially expelled. So, hydrophobic amino acids are more likely to favor formation of such an interface. But, this argument applies to both self and pathogen-derived peptides. Therefore, statistically, TCRs that bind more avidly to self-peptides should also bind more avidly to pathogen-derived peptides. There is some experimental evidence supporting this prediction (Mandl *et al*., 2013). Based on these considerations, we include the term, *R*(*s*, self), in Eq 1, which is the biochemical sequence similarity of peptide *s* to peptides derived from the human proteome. These peptides are also gathered from IEDB database (Vita *et al*., 2019) (Methods). Similar to Eq. 2, *R*(*s*, self) is defined as the number of self peptides whose alignment score with *s* is larger than a threshold value, *a*_*self*_ :

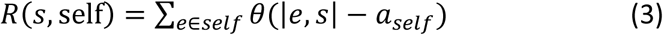

Now, *e* denotes a self-peptide.

We will use Eqs 1 - 3 to train a predictor of the immunodominance hierarchy of peptides targeted by CTLs in humans with different HLA alleles. The two parameters in our model are *a*_*self*_ and *a*_*pathogen*_, which will be determined by fitting the model to the training data (Methods). Given experimental measurements on the immunodominance hierarchy of peptides derived from pathogens in humans with different HLA alleles, we can constructed a binary classifier based on A(s, M). A peptide with *A*(*s, M*) larger than a threshold is classified as dominant and the others as nondominant. We trained and tested our model for *A*(*s, M*) as a predictor of immunodominance using experimental data on HIV peptides targeted by humans with different HLA alleles (Methods).

We systematically assembled data on HIV-1 specific CD8 T-cell responses, as determined by gamma interferon (IFN-γ) enzyme-linked immunospot (ELISPOT) assay, against a panel of up to 457 peptides including previously described optimal HIV-1 epitopes as defined in the Los Alamos National Laboratory HIV epitope database (www.hiv.lanl.gov) and epitope variants (Methods). Data was available from multiple cohorts of HIV-1 infected individuals at different stages of the infection and subsets of the data have been reported previously (Streeck *et al*., 2009; Pereyra *et al*., 2014). In total, optimal epitope specific CD8 T-cell data was available from 1102 individuals, including 619 individuals during acute and early infection, and 483 individuals during chronic infection of which 321 were considered spontaneous HIV controllers with median plasma HIV RNA levels ≤ 2,000 copies/ml. For the majority of individuals, the peptides used for T-cell response assessment were selected based on the individual’s HLA class I genotype. However, 314 Individuals with chronic infection had been tested against 267 optimal epitopes, irrespective of the individual’s HLA class I alleles. An average of 7 (range, 0-42) epitope specific CD8 T-cell responses were detected in the primary-infection cohort, while HIV-1-specific CD8 T-cell responses against an average of 20 epitopes (range, 0 to 95) were detected in chronically infected individuals. For our analysis, HLA class I restricted CD8 T-cell responses were considered only if the respective HLA allele was shared by at least 20 individuals in the data set. Table S1 in the supplemental material summarizes the frequencies of recognition for tested HIV-1-specific CD8 T-cell epitopes in the respective cohorts.

We used the percentage of patients with a given HLA allele responding to a given HIV peptide *s*, denoted as *p*(*s, M*), as the metric of immunodominance. The peptides which elicit response in more than 25% of tested patients with a given HLA allele were labelled as dominant and the others as non-dominant. Repeated 10-fold Cross-Validation was performed to train and test the model (Methods).

The performance of the *A*(*s, M*)-based classifier on the test sets is summarized as Receiver Operator Characteristic (ROC) curves (Fig 1). For the HIV acute infection group, the classifier has an Area Under the Curve (AUC) score of approximately 0.71 for the ROC curve. For the chronic infection group, the classifier has an AUC score of approximately 0.66. The superior performance in the acute infection dataset can be explained by the fact that, as HIV infection progresses, the virus mutates to escape CTL response, and as a result less immunodominant peptides are targeted by CTLs in the chronic phase (Streeck *et al*., 2009). The performance of the current model is compared to a T cell epitope immunogenicity prediction model developed by Calis et al (Calis *et al*., 2013), which is publicly available in IEDB. Our model shows superior performance as measured by the AUC (0.71 vs 0.57 for the acute group, 0.66 vs 0.34 for the chronic group; Fig 1). We also evaluated the importance of each of the three terms of *A*(*s, M*), the binding term, the term representing similarity to pathogenic peptides, and that representing similarity to human peptides by constructing partial models with one or two terms removed from *A*(*s, M*). The same training and testing procedures were repeated for these partial models. For both the acute and the chronic groups the partial models show less predictive power compared to the full model (Supplementary Fig. S1 and S2).

**Fig. 1.**
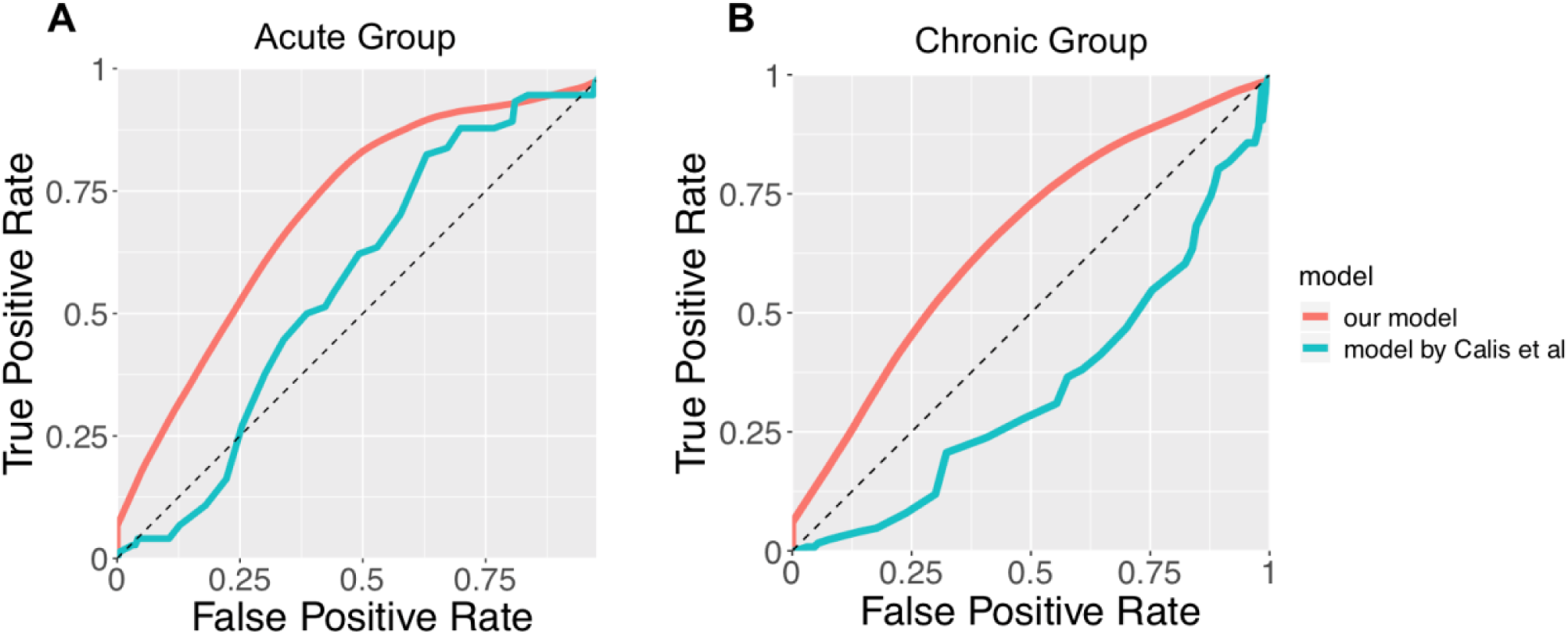
The ROC curve of the binary classifier based on our model (red), compared with the model developed by Calis et al (Calis *et al*., 2013) (green). (A) shows the ROC curves for the acute HIV infection group. The AUC of the red curve is 0.71. The AUC of the green curve is 0.57. (B) shows the ROC curves for the chronic HIV infection group. The AUC of the red curve is 0.66. The AUC of the green curve is 0.34.

### Only a fraction of SARS-CoV-2 peptides that bind to HLA molecules are immunogenic

Many research groups have identified peptides derived from SARS-CoV-2 that can bind with HLA molecules (Ahmed, Quadeer and McKay, 2020; Campbell *et al*., 2020; Grifoni *et al*., 2020; Nerli and Sgourakis, 2020; Prachar *et al*., 2020). Two different approaches were employed. In one approach, peptides that bind to different HLA molecules were identified based on experimental assays (Ahmed, Quadeer and McKay, 2020; Prachar *et al*., 2020). In the other approach, bioinformatic tools were used to identify peptides that bind to HLA molecules (Campbell *et al*., 2020; Grifoni *et al*., 2020; Nerli and Sgourakis, 2020). We used our trained classifier to predict the immunogenicity of peptides that were determined to bind to different HLA molecules experimentally, as reported by Ahmed et al (Ahmed, Quadeer and McKay, 2020) and Prachar et al (Prachar *et al*., 2020). Our classifier can be easily applied to the peptides reported by other groups too.

Ahmed et al (Ahmed, Quadeer and McKay, 2020) identified 187 SARS-CoV peptides that were suggested by HLA binding assays to bind to diverse HLA Class I molecules, and were identical in SARS-CoV-2. We further screened these peptides using our classifier (Methods), and found that only 74 of them are predicted to be immunogenic (Table 1 and Supplementary Table S2). These predicted peptides are associated with 33 different HLA alleles. Standard methods predict that this would enable coverage of 98.8% of the global population (i.e., 98.8% of the global population has at least one of these alleles), and 99.2% of US population (Methods).

**Table 1.**
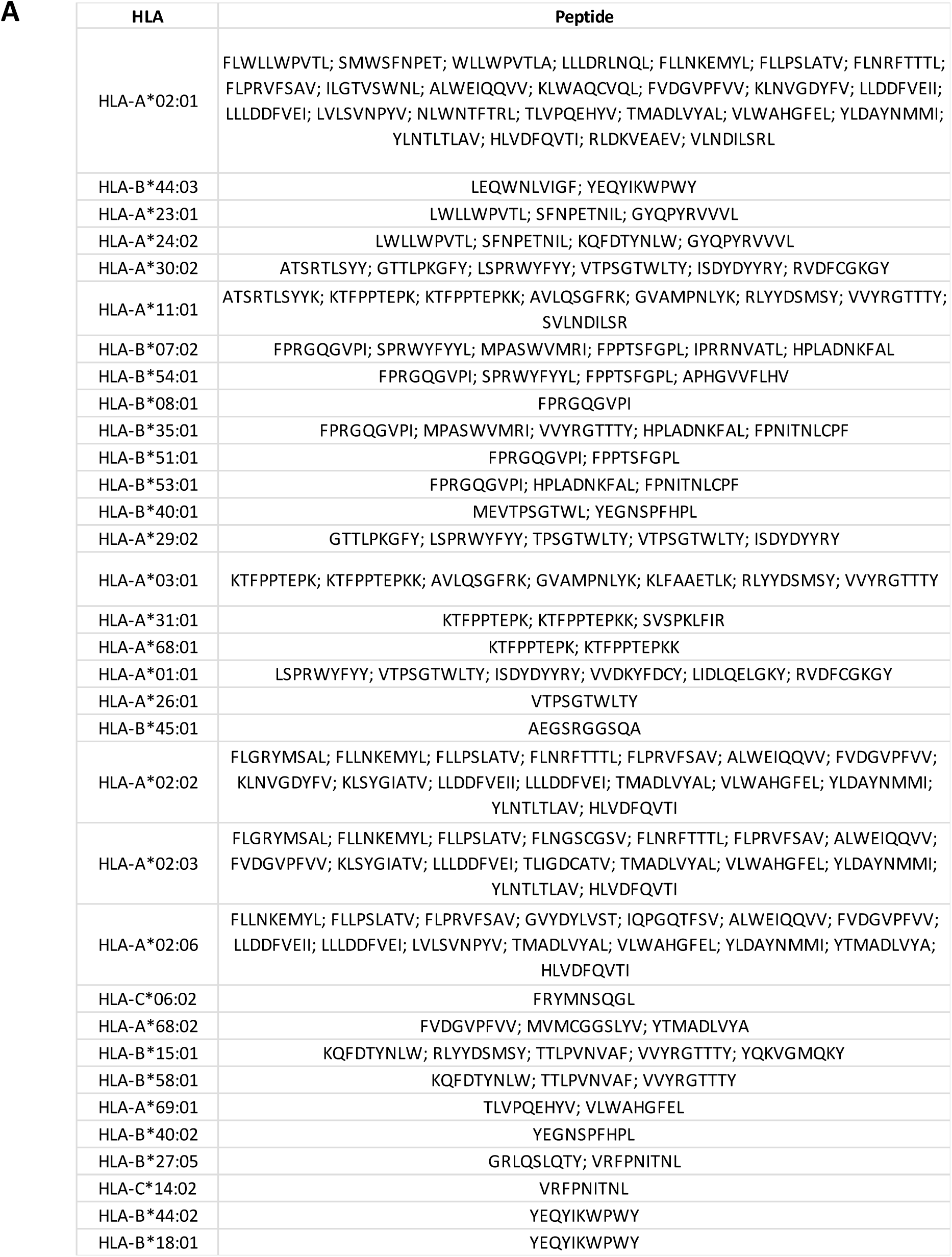
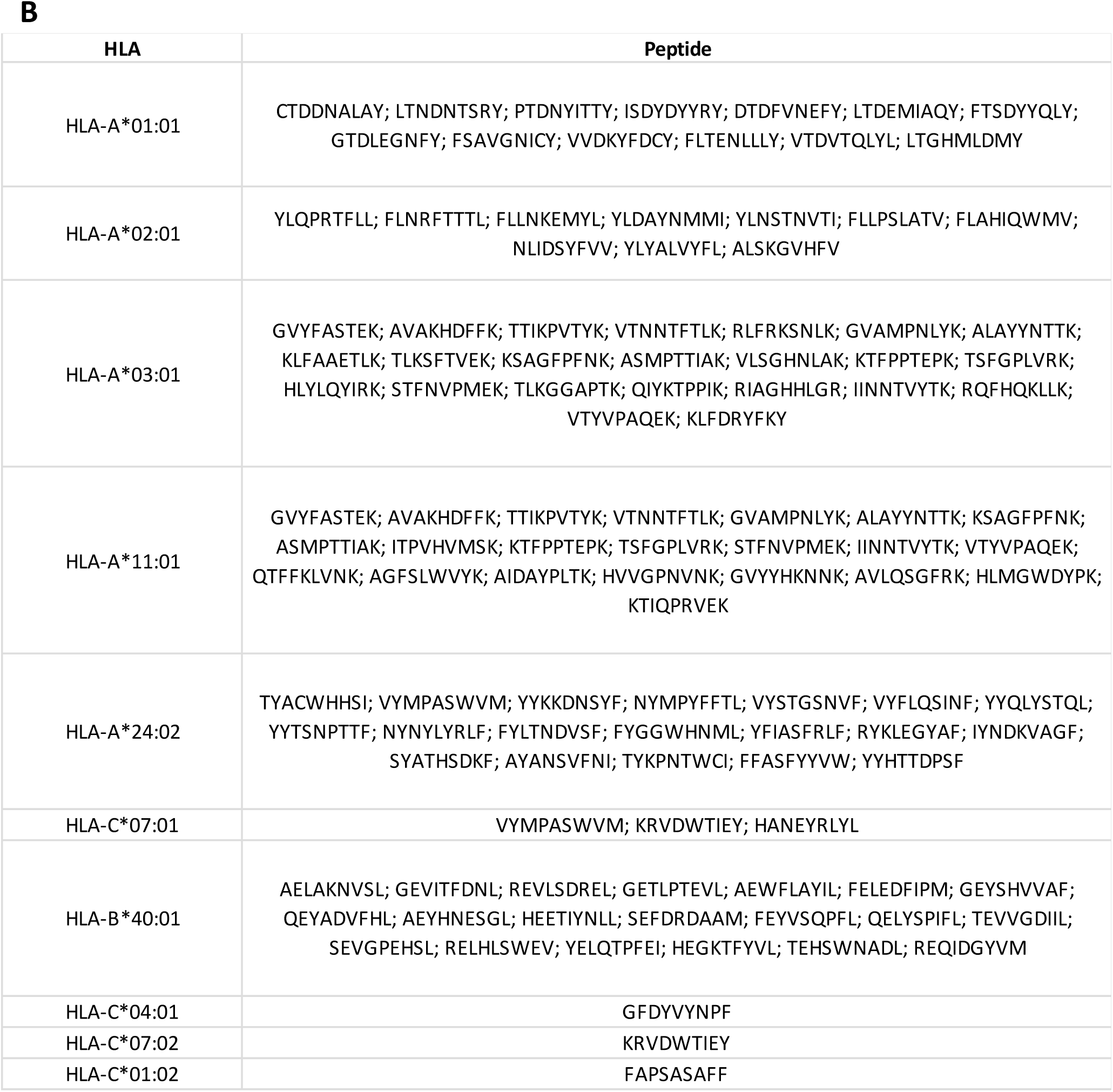
The top most immunogenic SARS-CoV-2 peptides predicted by our model. (A) shows the immunogenic peptides that are filtered from the peptide pool reported by Ahmed et al (Ahmed, Quadeer and McKay, 2020). (B) shows the immunogenic peptides that are filtered from those reported by Prachar et al (Prachar *et al*., 2020).

The same analysis was performed for the 152 SARS-CoV-2 peptides identified by Prachar et al (Prachar *et al*., 2020), which are also verified by HLA binding assays to be strong binders to diverse HLA Class I molecules. Our classifier predicted that 98 of them are immunogenic (Table 1 and Supplementary Table S2). They are associated with 10 different HLA alleles, which cover 94% of the global population and 93.2% of US population. These two sets of predicted immunogenic peptides can be combined together, which gives a total of 162 immunogenic peptides associated with 37 different HLA alleles (Supplementary Table S2). These HLA alleles can cover 99.6% of the global population and 99.7% of US population. On average each HLA allele is associated with 7.3 immunogenic peptides. Recall that the immunogenic peptides predicted by our model are defined as those that elicit a response in more than 25% of population with the associated HLA allele. With more than 7 immunogenic peptides associated with each HLA allele, it is likely that immunogenic CTL responses will be present in most people with the corresponding allele.

### CTL epitopes contained in the spike protein may not elicit sufficiently broad T-cell responses

Currently most SARS-CoV-2 vaccines only contain the spike protein of the virus as the immunogen (Akst, 2020). Thus, we wanted to test whether the immunogenic peptides from the spike protein alone can elicit CTL responses in a large portion of the population. Among the combined set of 162 predicted immunogenic peptides that we identified, 22 belong to the spike protein of the virus, and they are associated with 16 HLA alleles (Supplementary Table S2). These 16 HLA alleles cover 92.3% of the global population and 93.5% of the US population. However, on average each HLA allele is only associated with 1.8 immunogenic peptides. This relatively low number indicates that it is likely that most people with a particular allele will not mount immunogenic CTL responses. Therefore, including various viral proteins in the vaccine immunogen may be necessary in order to achieve broad coverage of CTL responses in a given population.

### There is significant overlap between immunogenic CTL epitopes in SARS-CoV-2 and common less pathogenic human coronaviruses

Unlike SARS-CoV-2 that causes severe respiratory disease, other less pathogenic coronaviruses circulating in the human population usually only cause mild diseases (like the common cold). Four common human coronavirus (HCoV), HCoV-229E(NC_002645.1), NL63(NC_005831.2), OC43(NC_006213.1) and HKU1(NC_006577.2) are responsible for 10-30% of upper respiratory tract infections in adults (Paules, Marston and Fauci, 2020). Given that memory T cell responses are likely induced in at least a fraction of the human population infected by these coronaviruses, we wanted to explore whether such memory responses could theoretically be induced/expanded following infection with SARS-CoV-2. Thus, we employed our classifier to identify common immunogenic peptide epitopes between HCoV and SARS-CoV-2. We first gathered a set of 38 HLA class I alleles that represent more than 99% of the world. We then applied our classifier to all possible overlapping 8-11mers in the proteome of SARS-CoV-2 and determined that there are 2311 immunogenic CTL epitopes associated with those 38 HLA alleles. We then further determined the unique set of immunogenic peptides that were common between SARS-CoV-2 and the four common HCoV. We found 46 shared immunogenic peptides, which are associated with 31 HLA alleles (Supplementary Table S3). These HLA alleles cover 98.6% of the global population and 99.0% of US population. On average each of these alleles are associated with 5.6 immunogenic peptides. Given this level of overlap between immunogenic epitopes between HCoVs and SARS-CoV-2, one can hypothesize that CTL memory responses elicited by past infection with common coronaviruses could respond to SARS-CoV-2 infection. We also accounted for the fact that each individual may have been infected by only a subset of the four less pathogenic coronaviruses. So, to determine a lower bound, we determined the immunogenic epitopes that are shared between SARS-CoV-2 and each of the four less pathogenic HCoV. The results are presented in Table 2. On average, 19 epitopes are common with any one of the less pathogenic human coronaviruses. The results shown in Table 2 lead us to suggest that some fraction of the human population has memory T cell responses that may target immunogenic SARS-CoV-2 CTL epitopes.

**Table 2.** Shared immunogenic peptides between SARS-CoV-2 and four common low pathogenicity human coronaviruses. The first column shows the number of shared immunogenic peptides between SARS-CoV-2 and each of the four viruses. The second column shows the number of HLA alleles associated with those peptides. The third and fourth column show the population coverage of those HLAs for the US and World, respectively. The fifth column shows the average number of immunogenic peptides associated with each HLA. The last row of the table shows the average of all these quantities.

|  | **N_peptide** | **N_HLA** | **HLA_coverage(US)** | **HLA_coverage (world)** | **Npep_per_HLA** |
| --- | --- | --- | --- | --- | --- |
| in229E | 8 | 24 | 93.60% | 90.20% | 1.67 |
| inHKU1 | 31 | 29 | 97.80% | 96.60% | 5.24 |
| inHL63 | 17 | 28 | 96.70% | 95.00% | 2.86 |
| inOC43 | 20 | 25 | 97.50% | 96.30% | 2.32 |
| **average** | **19** | **26.5** | **96.40%** | **94.53%** | **3.0225** |

## Discussion

In this work we developed a physics-based learning algorithm that predicts the CTL immunogenicity of peptides in humans with particular HLA alleles. This model was trained against a large experimental data set of clinical samples from HIV-infected individuals. When tested against experimental data for HIV epitopes, the model shows significantly improved performance compared to publicly available models (Calis *et al*., 2013).

Many groups have identified SARS-CoV-2 peptides that are able to bind with HLA molecules (Ahmed, Quadeer and McKay, 2020; Campbell *et al*., 2020; Grifoni *et al*., 2020; Nerli and Sgourakis, 2020; Prachar *et al*., 2020), either using experimental assays or bioinformatic tools. We screened these peptides for immunogenicity using our algorithm. Specifically, we studied the peptides suggested by Ahmed et al (Ahmed, Quadeer and McKay, 2020) and Prachar et al (Prachar *et al*., 2020), but our model can be applied to filter peptides suggested by other groups. Our results suggest that only a fraction of the peptides that bind to HLA molecules are likely to be immunogenic. But, the combined set of SARS-CoV-2 peptides that we predict to be immunogenic among known HLA binders provides broad coverage of the global population. These predictions need to be further experimentally validated, but we hope that our results can guide the choice of peptide – HLA combinations that need to be tested experimentally for determining the CTL immunogenicity of SARS-CoV-2 peptides in humans.

We note that mutations in SARS-CoV-2 thus far are uncommon (Ahmed, Quadeer and McKay, 2020), and our results suggest that the immunogenic peptide epitopes in the virus’ proteome may be sufficient to provide broad coverage of the population. Therefore, determination of mutational vulnerabilities of the virus to focus CTL responses to special epitopes, as has been done for HIV (Létourneau *et al*., 2007; Dahirel *et al*., 2011; Shekhar *et al*., 2013; Ferguson *et al*., 2013; Hayton *et al*., 2014; Mann *et al*., 2014; Abdul-Jawad *et al*., 2016; Barton *et al*., 2016; Louie *et al*., 2018; Ahmed *et al*., 2019; Gaiha *et al*., 2019), is likely not necessary. Whole proteome immunogens should suffice in a vaccine that aims to elicit potent CTL responses that provides broad population coverage.

Since most SARS-CoV-2 vaccines in development use only the spike protein as immunogen, we also analyzed whether peptides from the spike alone can yield broad CTL coverage over the global population. Based on our analysis, the immunogenic spike peptides alone are unlikely to provide such broad coverage from the standpoint of CTL responses. Therefore, to get broad CTL coverage, an immunogen consisting of other SARS-CoV-2 proteins might be necessary. This is potentially significant if antibody responses to SARS-CoV-2 prove not to be durable, as reported for SARS-CoV.

With regards to common human coronaviruses which have likely infected substantially more individuals than SARS-CoV-2 despite the current pandemic, we predict that there is overlap between the immunogenic CTL epitopes among these viruses. This suggests that memory T cells directed against less pathogenic coronaviruses could target immunogenic SARS-CoV-2 epitopes upon infection or following vaccination with a SARS-CoV-2 immunogen. Clinical outcomes and the course of disease during SARS-CoV-2 infection are extremely heterogeneous ranging from asymptomatic disease to death (Fu *et al*., 2020). Whether pre-exisiting HCoV specific memory T-cells actually play a disease modifying or even protective role needs to be determined. The common immunogenic epitopes that we have identified could serve as a first target that might be examined.

Although validated by HIV CTL response data, this model is not yet experimentally tested in the case of SARS-CoV-2 or other viral infections. It is important to further validate, and potentially elaborate, the model by testing against experimental data for diverse viruses. More data will also help improve the model. Currently the model contains two parameters *a*_*self*_ and *a_pathogen_*, which are the cutoff thresholds for similarity to self and pathogenic peptides, respectively. These two parameters are used for all HLA alleles. However, it is known that peptides bound to different alleles can use different peptide residues to make primary contacts with the TCR. So, a model with allele-specific similarity cutoff thresholds might improve the performance. This will require training our model against more extensive datasets. Since short peptides derived from the proteome do not carry long-range information about the pathogen, if our model is further validated and elaborated, it may be a useful and simple tool for rapid identification of immunogenic CTL epitopes contained in diverse new endemic-causing pathogens that will undoubtedly emerge in the future.

## Methods

### Training of the model

The model is trained and tested by repeated 10-fold cross-validation. For the 10-fold cross-validation, the dataset is evenly divided into 10 subsets. One subset is taken as the test set and the remaining 9 subsets as the training set. The two parameters of the model, *a*_*self*_ and *a*_*pathogen*_, are determined by maximizing the Spearman correlation coefficient between the CTL response metric *A*(*s, M*) and the response percent *p*(*s, M*) for the training set. Then the trained *A*(*s, M*) is evaluated on the test set. We repeated the 10-fold cross validation procedure 20 times, and the dataset is reshuffled before each repetition. The Spearman correlation coefficient between *A*(*s, M*) and *p*(*s, M*) on the test sets are summarized in Supplementary Fig. S1.

The binary classifier based on *A*(*s, M*) is constructed in the following way: the peptide with *A*(*s, M*) larger than a threshold is classified as dominant and the others as nondominant. When the classifier is used to predict immunodominant SARS-Cov-2 peptides, the threshold is chosen to be 7024. At this threshold the True Positive Rate and the True Negative Rate of the classifier are equal for the acute HIV infection patient data.

### Self and Foreign peptides

The foreign peptides are retrieved from the Immune Epitope Database (IEDB) (Vita *et al*., 2019) using the following search criteria: “Epitope: Linear; Assay: T Positive; MHC Restriction: MHC I; Disease: Infectious disease; Host: Human”. To avoid bias in training and testing the model, we excluded HIV peptides from the retrieved foreign peptides. The self peptides are retrieved from the IEDB database using the following search criteria: “Epitope: Linear; Assay: MHC Ligand Assay Positive; Organism: Homo Sapiens”.

### Population coverage

The population coverage of the HLA alleles is estimated using the publicly available tool on IEDB (Bui *et al*., 2006) (http://tools.iedb.org/population/).

### Experimental methods

Data of HIV-specific CD8 T-cell responses from a total of 1102 HIV-1-infected individuals were included in this analysis, consisting of previously published and unpublished data (Streeck *et al*., 2009; Pereyra *et al*., 2014). 619 Individuals from primary-infection cohorts in North America, Germany, and Australia were included. 483 Individuals with chronic HIV infection recruited from outpatient clinics at local Boston hospitals and referred from providers throughout the United States were included. 321 of the chronic infection cohort were classified as spontaneous HIV controllers as their median plasma HIV RNA levels were ≤2,000 copies/ml. The study was approved by the respective institutional review boards and was conducted in accordance with the human experimentation guidelines of Massachusetts General Hospital.

### HLA typing

High-resolution class I HLA typing for HLA-A, -B, and -C was performed at the National Cancer Institute, National Institutes of Health, Frederick, MD, using sequence-based typing protocols developed by the 13th International Histocompatibility Workshop (Hansen, 2005) on DNA that was extracted from whole blood.

### Assessment of HIV-specific CD8+ T-cell responses

HIV-1-specific CD8 T-cell responses were quantified by gamma interferon (IFN-γ) enzyme-linked immunospot (ELISPOT) assay on previously cryopreserved peripheral blood mononuclear cells (PBMC), using a panel of up to 457 peptides including previously described optimal HIV-1 epitopes (www.hiv.lanl.gov) and epitope variants, as previously described (Streeck *et al*., 2009; Pereyra *et al*., 2014). Medium alone served as a negative control and phytohemagglutinin (PHA) as a positive control. The numbers of specific IFN-γ-secreting T cells were enumerated using an automated ELISPOT reader (Cellular Technology Ltd., Shaker Heights, OH) and expressed as spot-forming cells (SFCs)/10^6^ PBMC. A response was considered positive only if there were ≥ 55 SFCs/10^6^ PBMC and SFC/10^6^ PBMC was at least three times greater than mean background of the SFC/10^6^ PBMC in the negative wells.

## Acknowledgments

This work was supported by the National Institutes of Health (AI138790 to B.J.), NSF grant # PHY-2026995 to A.G. and A.K.C. A.G., B.J. and A.K.C.were supported by the Ragon Institute of MGH, MIT, & Harvard. This project has been funded in whole or in part with federal funds from the Frederick National Laboratory for Cancer Research, under Contract No. HHSN261200800001E. The content of this publication does not necessarily reflect the views or policies of the Department of Health and Human Services, nor does mention of trade names, commercial products, or organizations imply endorsement by the U.S. Government. This Research was supported in part by the Intramural Research Program of the NIH, Frederick National Lab, Center for Cancer Research.

