## Supplementary Figures and Tables for "Predicting the Immunogenicity of T cell epitopes: From HIV to SARS-CoV-2"

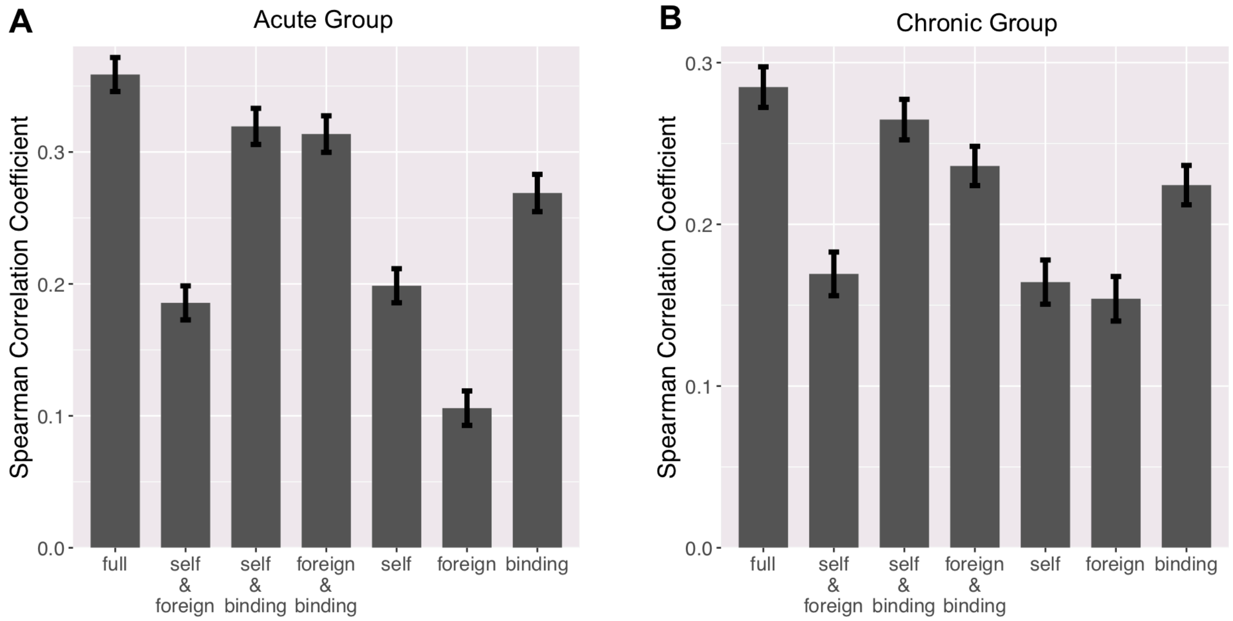


Fig. S1. The spearman correlation coefficient between $A(s, M)$ and $p(s, M)$ on the test sets. The full model is compared with partial models where one or two of the three terms of $A(s, M)$ are missing. (A) corresponds to the acute group. (B) corresponds to the chronic group.

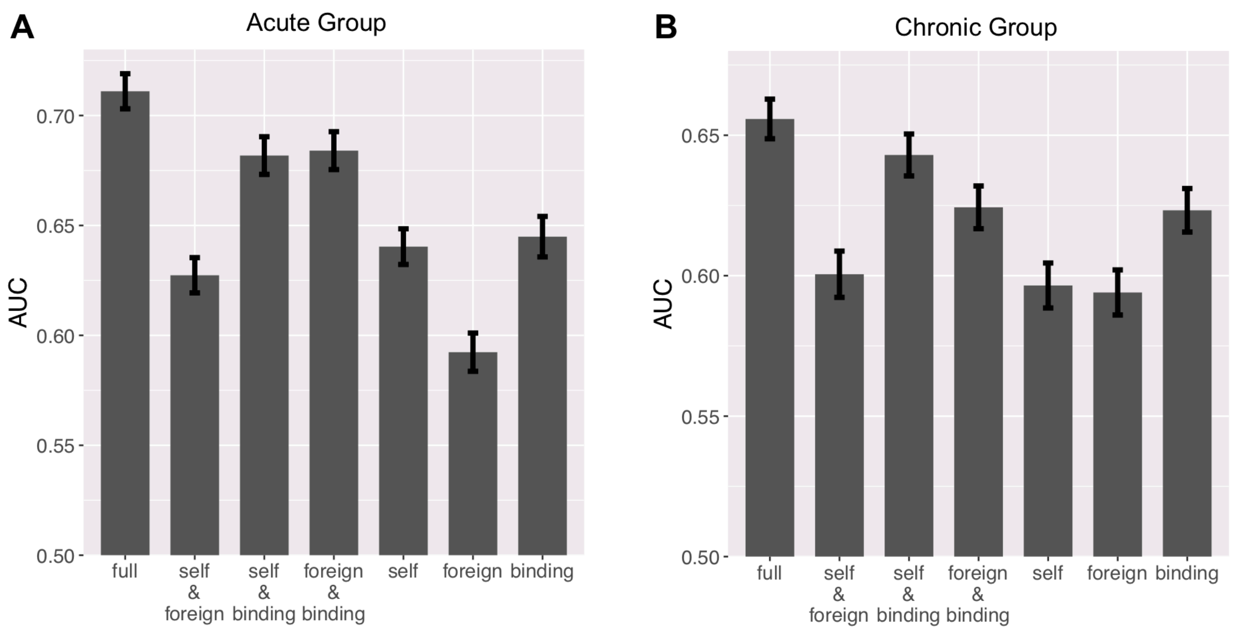


Fig. S2. The AUC of the binary classifier based on our model. The full model is compared with partial models where one or two of the three terms of $A(s, M)$ are missing. (A) corresponds to the acute group. (B) corresponds to the chronic group.

Table S1. The percentage of acutely/chronically infected patients responding to optimally defined HLA class I matched HIV peptides.
(Supplementary_Table_S1.xlsx)

Table S2. The full list of predicted immunogenic SARS-Cov-2 peptides which are filtered from datasets by Ahmed et al [1] and by Prachar et al [2].
(Supplementary_Table_S2.xlsx)

Table S3. The list of predicted immunogenic peptides that are shared between SARS-Cov-2 peptides and four common coronaviruses.
(Supplementary_Table_S3.xlsx)

#### Supplementary Table S1

| Acute Infection | | | | Chronic Infection | | | |
|---|---|---|---|---|---|---|---|
| HLA | epitope name | epitope | response percent | HLA | epitope name | epitope | response percent |
| A25:01 | EW10 | ETINEEAAEW | 0.826086957 | B57:03 | KF11 | KAFSPEVIPMF | 0.921568627 |
| B27:05 | KK10 | KRWIILGLNK | 0.805555556 | B51:01 | LI9 | LPPVVAKEI | 0.857142857 |
| B08:01 | FL8 | FLKEKGGL | 0.720338983 | B57:01 | KF11 | KAFSPEVIPMF | 0.841463415 |
| B57:01 | TW10 | TSTLQEQIGW | 0.72 | B08:01 | EI8 | EIYKRWII | 0.840909091 |
| B08:01 | FL8_K3Q | FLQEKGGL | 0.714285714 | A25:01 | EW10 | ETINEEAAEW | 0.827586207 |
| A24:02 | RW8_Y2F | RFPLTFGW | 0.681818182 | B15:01 | GY9 | GLNKIVRMY | 0.794117647 |
| A03:01 | RK9 | RLRPGGKKK | 0.659574468 | B57:03 | IW9 | ISPRTLNAW | 0.764705882 |
| B40:01 | KL9 | KEKGGLEGL | 0.646341463 | A29:02 | FY9 | FNCGGEFFY | 0.756756757 |
| B14:02 | EL9 | ERYLKDQQL | 0.631578947 | B57:01 | IW9 | ISPRTLNAW | 0.743902439 |
| B08:01 | FL8_K5Q | FLKEQGGL | 0.619047619 | B14:02 | DA9 | DRFYKTLRA | 0.733333333 |
| A03:01 | RY10 | RLRPGGKKKY | 0.602836879 | A68:02 | GA10 | GAETFYVDGA | 0.724137931 |
| B51:01 | LI9 | LPPVVAKEI | 0.59375 | B44:03 | AW11_D7E | AEQASQEVKNW | 0.720930233 |
| B14:02 | DA9 | DRFYKTLRA | 0.58974359 | A30:02 | KYY9 | KIQNFRVYY | 0.711111111 |
| A32:01 | RW9 | RIKQIINMW | 0.580645161 | B53:01 | QW9 | QASQEVKNW | 0.710526316 |
| B40:02 | KL9 | KEKGGLEGL | 0.56 | A25:01 | QW11 | QAISPRTLNAW | 0.7 |
| B08:01 | EI8 | EIYKRWII | 0.559322034 | B51:01 | RI8 | RPLVTIKI | 0.689655172 |
| A24:02 | RW8 | RYPLTFGW | 0.551020408 | B08:01 | FL8 | FLKEKGGL | 0.681818182 |
| B57:01 | IW9 | IVLPEKDSW | 0.52 | A29:02 | SY10 | SFNCGGEFFY | 0.675675676 |
| B51:01 | TI8 | TAFTIPSI | 0.515625 | A29:02 | YT9 | YFPDWQNYT | 0.675675676 |
| B27:05 | VL9 | VRHFPRIWL | 0.515151515 | A26:01 | EL9 | EVIPMFSAL | 0.633333333 |
| B44:02 | AW11_S5T | AEQATQDVKNW | 0.5 | B53:01 | YF9 | YPLTFGWCF | 0.631578947 |
| C08:02 | AL9 | AAVDLSHFL | 0.490196078 | B35:01 | VY8 | VPLRPMTY | 0.630434783 |
| A33:01 | ER9 | EYRKILRQR | 0.454545455 | A30:02 | RY11 | RSLYNTVATLY | 0.622222222 |
| C07:01 | KY11_K1Q | QRQEILDLWVY | 0.452380952 | B57:01 | TW10_G9A | TSTLQEQIAW | 0.62 |
| A29:02 | SY9 | SFEPIPIHY | 0.45 | B53:01 | YY9 | YPLTFGWCY | 0.605263158 |
| B08:01 | RL9 | RQGLERALL | 0.449152542 | A03:01 | RK9 | RLRPGGKKK | 0.594059406 |
| A03:01 | KK9 | KIRLRPGGK | 0.446808511 | C08:02 | AL9 | AAVDLSHFL | 0.58974359 |
| A11:01 | ALK9 | AVDLSHFLK | 0.445945946 | A03:01 | QK10 | QVPLRPMTYK | 0.584158416 |
| B57:01 | HQ10 | HTQGYFPDWQ | 0.441860465 | B44:02 | AW11_D7E | AEQASQEVKNW | 0.576271186 |
| A26:01 | EL9 | EVIPMFSAL | 0.4375 | B40:01 | KL9 | KEKGGLEGL | 0.571428571 |
| A24:02 | RW8_F6L | RYPLTLGW | 0.428571429 | A03:01 | RY10 | RLRPGGKKKY | 0.57 |
| B08:01 | RL9_L9Q | RQGLERALQ | 0.428571429 | A11:01 | AK11 | ACQGVGGPGHK | 0.555555556 |
| A29:02 | SY10 | SFNCGGEFFY | 0.425 | C05:01 | AM12 | AEQASQEVKNWM | 0.55 |
| A11:01 | AK11 | ACQGVGGPGHK | 0.418918919 | B52:01 | RI8 | RMYSPTSI | 0.55 |
| A29:02 | FY9 | FNCGGEFFY | 0.411764706 | A32:01 | PW10 | PIQKETWETW | 0.545454545 |
| A29:02 | YT9 | YFPDWQNYT | 0.411764706 | A03:01 | KK9 | KIRLRPGGK | 0.544554455 |
| B07:02 | IL9 | IPRRIRQGL | 0.411290323 | B57:01 | QW9 | QASQEVKNW | 0.543209877 |
| B40:01 | KSL9 | KELYPLTSL | 0.409638554 | A11:01 | TK8 | TLYCVHQK | 0.52173913 |
| A32:01 | PW10 | PIQKETWETW | 0.40625 | B18:01 | YY9 | YPLTFGWCY | 0.515151515 |
| B57:01 | KF11 | KAFSPEVIPMF | 0.4 | B14:02 | EL9 | ERYLKDQQL | 0.5 |
| B07:02 | GL9_G3A | GPAHKARVL | 0.392857143 | B57:03 | TW10_G9A | TSTLQEQIAW | 0.5 |
| A11:01 | QK10 | QVPLRPMTYK | 0.391891892 | B40:01 | QL10 | QELKNSAVSL | 0.5 |
| A25:01 | QW11 | QAISPRTLNAW | 0.391304348 | B07:02 | TL10 | TPGPGVRYPL | 0.490566038 |
| C07:01 | KY11 | KRQEILDLWVY | 0.390804598 | B57:03 | FF9 | FSPEVIPMF | 0.490196078 |
| A24:02 | KW9 | KYKLKHIVW | 0.387755102 | C03:04 | AL9 | AFHHVAREL | 0.485714286 |
| A30:01 | KYY9 | KIQNFRVYY | 0.382352941 | A68:02 | EV10 | ETYGDTWTGV | 0.482758621 |
| C12:03 | CC8 | CCFHCQVC | 0.380952381 | B35:01 | TY9 | TVLDVGDAY | 0.47826087 |
| C07:01 | KY11_V10I | KRQEILDLWIY | 0.375 | A02:01 | SL9 | SLYNTVATL | 0.477777778 |
| A03:01 | QK10 | QVPLRPMTYK | 0.368794326 | B40:01 | IL8 | IETVPVKL | 0.476190476 |
| B44:02 | AW11 | AEQASQDVKNW | 0.36809816 | B53:01 | EW10 | EPVDPRLEPW | 0.473684211 |
| B57:01 | HW9 | HTQGYFPDW | 0.367346939 | B07:02 | SM9 | SPAIFQSSM | 0.471698113 |
| B44:02 | EW9 | EEMNLPGRW | 0.360294118 | B35:01 | HY9 | HPDIVIYQY | 0.466666667 |
| B51:01 | TI8_I8T | TAFTIPST | 0.357142857 | A30:02 | KIY9 | KLNWASQIY | 0.466666667 |
| B40:02 | AL9_A1G | GALDLSHFL | 0.35 | A02:01 | FK10 | FLGKIWPSYK | 0.463687151 |
| B44:02 | AW11_D7E | AEQASQEVKNW | 0.347826087 | B57:01 | YT9 | YFPDWQNYT | 0.463414634 |
| C03:04 | AL9_A1G | GALDLSHFL | 0.346153846 | A29:02 | LY9 | LYNTVATLY | 0.459459459 |
| A31:01 | RR11 | RLRDLLLIVTR | 0.346153846 | C04:01 | SF9 | SFNCGGEFF | 0.458715596 |
| B40:01 | IL10 | IEIKDTKEAL | 0.325301205 | B35:01 | VY10 | VPLDEDFRKY | 0.456521739 |
| C05:01 | AM12 | AEQASQEVKNWM | 0.317647059 | B57:03 | TW10 | TSTLQEQIGW | 0.450980392 |
| B15:01 | GY9 | GLNKIVRMY | 0.315789474 | C07:01 | KY11 | KRQEILDLWVY | 0.45045045 |
| B51:01 | EI9 | EAVRHFPRI | 0.3125 | A31:01 | RR11 | RLRDLLLIVTR | 0.448275862 |
| C03:04 | AL9 | AAVDLSHFL | 0.304964539 | B57:01 | HW9 | HTQGYFPDW | 0.444444444 |
| B07:02 | GL9 | GPGHKARVL | 0.301587302 | B15:01 | IY10 | ILKEPVHGVY | 0.441176471 |

| | | | | | | | |
|---|---|---|---|---|---|---|---|
| B57:01 | IW9 | ISPRTLNAW | 0.3 | B40:01 | IL10 | IEIKDTKEAL | 0.434782609 |
| B57:01 | SW10 | STTVKAACWW | 0.3 | B35:01 | NQY9 | NPDIVIYQY | 0.434782609 |
| B07:02 | RL9_T4D | RPMDYKAAL | 0.3 | B57:03 | TW10 | TSNLQEQIAW | 0.433333333 |
| B35:01 | VY8 | VPLRPMTY | 0.279661017 | B57:01 | HQ10 | HTQGYFPDWQ | 0.432098765 |
| B07:02 | RY10 | RQDILDLWIY | 0.277777778 | B44:02 | EW9_E2D | EDMNLPGRW | 0.428571429 |
| B40:02 | AV9 | AEWDRVHPV | 0.271604938 | C06:02 | YT9 | YFPDWQNYT | 0.428571429 |
| A29:02 | LY9 | LYNTVATLY | 0.264705882 | B44:03 | EW9_E2D | EDMNLPGRW | 0.418604651 |
| B07:02 | RL9_T4N | RPMNYKAAL | 0.259259259 | B07:02 | IL9 | IPRRIRQGL | 0.41509434 |
| A68:01 | IR11 | ITKGLGISYGR | 0.257142857 | B57:01 | AW9 | AVRHFPRIW | 0.414634146 |
| B27:05 | RK10 | RRWIQLGLQK | 0.25 | A68:02 | IR11 | ITKGLGISYGR | 0.413793103 |
| B07:02 | TL10_V6I_R7K | TPGPGIKYPL | 0.25 | B57:03 | HQ10 | HTQGYFPDWQ | 0.411764706 |
| C07:01 | RY11 | RRQDILDLWIY | 0.242622951 | B40:01 | IL9 | IEELRQHLL | 0.409090909 |
| B07:02 | RL9 | RPMTYKAAL | 0.241071429 | A11:01 | QK10 | QVPLRPMTYK | 0.407407407 |
| B08:01 | WM8_E6D | WPTVRDRM | 0.238095238 | B14:02 | CC9 | CRAPRKKGC | 0.4 |
| B14:02 | SL9 | SAEPVPLQL | 0.236842105 | A26:01 | EY9 | ETKLGKAGY | 0.4 |
| B35:01 | VY8_Y8F | VPLRPMTF | 0.227272727 | B57:01 | TW10 | TSNLQEQIGW | 0.4 |
| A33:01 | TW9 | TRYPLTFGW | 0.227272727 | B57:03 | TW10 | TSNLQEQIGW | 0.4 |
| B27:05 | GY10 | GRRGWEALKY | 0.222222222 | C07:01 | RY11 | RPQVPLRPMTY | 0.396396396 |
| B27:05 | IK9 | IRLRPGGKK | 0.222222222 | B07:02 | GL9 | GPGHKARVL | 0.396226415 |
| B07:02 | GL9_G3S | GPSHKARVL | 0.222222222 | B57:03 | HW9 | HTQGYFPDW | 0.392156863 |
| C08:02 | SL9 | SAEPVPLQL | 0.22 | B57:01 | IW9 | IVLPEKDSW | 0.390243902 |
| A26:01 | EY9 | ETKLGKAGY | 0.21875 | C08:02 | TL9 | TPQDLNTML | 0.384615385 |
| A24:02 | RL11 | RDYVDRFFKTL | 0.214285714 | B57:01 | TW10 | TSNLQEQIAW | 0.38 |
| A03:01 | ALK9 | AVDLSHFLK | 0.212765957 | B57:01 | TW10 | TSTLQEQIGW | 0.37804878 |
| B35:01 | NQY9 | NPDIVIYQY | 0.211864407 | A30:02 | KY11 | KEKGGLEGLIY | 0.377777778 |
| B35:01 | VY10_V1Y_5EK | YPLDKDFRKY | 0.208333333 | A11:01 | ATK9 | AIFQSSMTK | 0.37037037 |
| C15:02 | RL9 | RAIEAQQHL | 0.206896552 | B57:01 | SW10 | STTVKAACWW | 0.37037037 |
| B40:01 | IL8 | IETVPVKL | 0.202702703 | C07:02 | RY11 | RPQVPLRPMTY | 0.365079365 |
| A02:01 | FK10 | FLGKIWPSYK | 0.198757764 | C08:02 | AL9 | AAVDLSHFL | 0.363636364 |
| C05:01 | SL9 | SAEPVPLQL | 0.196078431 | B07:02 | RY10 | RQDILDLWIY | 0.358490566 |
| A02:01 | SL9 | SLYNTVATL | 0.195652174 | B51:01 | EI9 | EAVRHFPRI | 0.355555556 |
| B27:05 | RI10 | RRQDILDLWI | 0.194444444 | B57:01 | KY10 | KAVRLIKFLY | 0.353658537 |
| B44:03 | LY9 | LYNTVATLY | 0.191489362 | B57:03 | SW10 | STTVKAACWW | 0.352941176 |
| B35:01 | VY8_T7D | VPLRPMDY | 0.19047619 | A23:01 | RL9 | RYLKDQQLL | 0.35 |
| B07:02 | TL10 | TPGPGVRYPL | 0.19047619 | A30:01 | RY11 | RSLYNTVATLY | 0.342857143 |
| A01:01 | GY9 | GSEELRSLY | 0.186335404 | C08:02 | HL9 | HLVWASREL | 0.342105263 |
| B57:01 | YY9 | YTPGPGIRY | 0.186046512 | A03:01 | ATK9 | AIFQSSMTK | 0.336633663 |
| B07:02 | RV9 | RPMTYKAAV | 0.182539683 | A11:01 | IK10 | IYQEPFKNLK | 0.333333333 |
| B57:01 | QW9 | QASQEVKNW | 0.18 | A30:02 | KY9 | KQNPDIVIY | 0.333333333 |
| A30:01 | KIY9 | KLNWASQIY | 0.176470588 | B07:02 | RL9 | RPMTYKAAL | 0.333333333 |
| B07:02 | SM9 | SPAIFQSSM | 0.174603175 | B51:01 | TI8 | TAFTIPSI | 0.333333333 |
| B18:01 | YY9 | YPLTFGWCY | 0.174603175 | A29:02 | SY9 | SFEPIPIHY | 0.324324324 |
| C04:01 | SF9 | SFNCGGEFF | 0.165517241 | B07:02 | RV9 | RPMTYKAAV | 0.320754717 |
| A11:01 | QVK9 | QIYAGIKVK | 0.162162162 | C15:02 | RL9 | RYLKDQQLL | 0.32 |
| B08:01 | WM8 | WPTVRERM | 0.161016949 | A02:01 | IV9 | ILKEPVHGV | 0.312849162 |
| A02:01 | GL9_F3V | GAVDLSFFL | 0.159090909 | C07:02 | KY11 | KRQEILDLWVY | 0.305084746 |
| B15:03 | FY10 | FQTKGLGISY | 0.157894737 | B08:01 | RL9 | RQGLERALL | 0.302325581 |
| C08:02 | TL9 | TPQDLNTML | 0.156862745 | A01:01 | YT9 | YFPDWQNYT | 0.301204819 |
| B40:01 | IL9 | IEELRQHLL | 0.156626506 | C14:02 | LL9 | LYNTVATL | 0.3 |
| A02:01 | GL9_F3M | GAMDLSFFL | 0.155555556 | A11:01 | ALK9 | AVDLSHFLK | 0.296296296 |
| B35:01 | DL9 | DPNPQEVVL | 0.152542373 | B15:01 | LY12 | LVGKLNWASQIY | 0.294117647 |
| C06:02 | YT9 | YFPDWQNYT | 0.150684932 | B57:01 | IF9 | ISKKAKGWF | 0.292682927 |
| A03:01 | QR9 | QIYPGIKVR | 0.14893617 | C03:04 | RL9 | RYLKDQQLL | 0.285714286 |
| B40:02 | RL9 | REPHNEWTL | 0.148148148 | C04:01 | QW9 | QASQEVKNW | 0.28440367 |
| B44:02 | LY9 | LYNTVATLY | 0.147727273 | B07:02 | RI10 | RPNNNTRKSI | 0.283018868 |
| B07:02 | RI10 | RPNNNTRKSI | 0.14516129 | C08:02 | RV9 | RAEQASQEV | 0.282051282 |
| B35:01 | VY10 | VPLDEDFRKY | 0.144067797 | B44:03 | LY9 | LYNTVATLY | 0.279069767 |
| A24:02 | RL9 | RYLKDQQLL | 0.142857143 | B57:03 | AW9 | AVRHFPRIW | 0.274509804 |
| B57:01 | IF9 | ISKKAKGWF | 0.14 | C03:03 | AL9 | AFHHVAREL | 0.272727273 |
| B57:01 | KY10 | KAVRLIKFLY | 0.14 | A32:01 | RW9 | RIKQIINMW | 0.272727273 |
| A02:01 | YI9 | YTAFTIPSI | 0.139751553 | B35:01 | WF9 | WASRELERF | 0.260869565 |
| A02:01 | SL9_A7V | SLYNTVVTL | 0.137254902 | A03:01 | AK10 | AVFIHNFKRK | 0.25 |
| A02:01 | AL9 | AIIRILQQL | 0.136645963 | B18:01 | LY10 | LADQLIHLHY | 0.25 |
| B35:01 | PY9 | PPIPVGDIY | 0.13559322 | A03:01 | RK10 | RIRTWKSLVK | 0.245098039 |
| B44:02 | AY9 | AENLWVTVY | 0.134969325 | A02:01 | AL9 | AIIRILQQL | 0.241573034 |
| A01:01 | RY9 | RRGWEVLKY | 0.12962963 | A11:01 | QKK9 | QIIEQLIKK | 0.240740741 |
| B39:01 | GL9 | GHQAAMQML | 0.129032258 | A11:01 | QVK9 | QIYAGIKVK | 0.240740741 |
| B07:02 | FL9 | FPRIWLHGL | 0.128 | A68:01 | IR11 | ITKGLGISYGR | 0.24 |

| | | | | | | | |
|---|---|---|---|---|---|---|---|
| B51:01 | EI9 | EKEGKISKI | 0.125 | A30:02 | LF11 | LVWASRELERF | 0.239130435 |
| A02:01 | GL9_F3L | GALDLSFFL | 0.125 | B40:01 | LS9 | LEKHGAITS | 0.238095238 |
| C03:04 | RL9 | RAIEAQQHL | 0.124031008 | B57:03 | YT9 | YFPDWQNYT | 0.235294118 |
| B40:02 | GI9 | GELDRWEKI | 0.12345679 | A03:01 | ALK9 | AVDLSHFLK | 0.232323232 |
| A68:01 | DR11 | DTWAGVEAIIR | 0.122807018 | A03:01 | RK11 | RMRGAHTNDVK | 0.23 |
| A68:01 | GA10 | GAETFYVDGA | 0.122807018 | A30:01 | LF11 | LVWASRELERF | 0.228571429 |
| A11:01 | AK9 | AIFQSSMTK | 0.121621622 | A11:01 | AK10 | AVFIHNFKRK | 0.226415094 |
| B07:02 | HI10 | HPRVSSEVHI | 0.120967742 | C01:02 | VL8 | VIPMFSAL | 0.225 |
| B35:01 | TY9 | TVLDVGDAY | 0.118644068 | A02:01 | PL10 | PLTFGWCYKL | 0.217877095 |
| C08:02 | RV9 | RAEQASQEV | 0.117647059 | B40:01 | KSL9 | KELYPLTSL | 0.217391304 |
| A30:01 | RY11 | RSLYNTVATLY | 0.117647059 | A30:02 | RV10 | RMRGAHTNDV | 0.209302326 |
| B08:01 | FL8_E4K | FLKKKGGL | 0.115384615 | B07:02 | HI10 | HPRVSSEVHI | 0.20754717 |
| A02:01 | RI9 | RILQQLLFI | 0.114906832 | B07:02 | TL9 | TPQDLNTML | 0.20754717 |
| B40:02 | TL8 | TERQANFL | 0.111111111 | B15:01 | TY11 | TQGYFPDWQNY | 0.205882353 |
| B51:01 | LI9 | LPCRIKQII | 0.109375 | B08:01 | GL9 | GPKVKQWPL | 0.204545455 |
| B39:01 | TL9 | TPQDLNTML | 0.107142857 | B51:01 | RL9 | RAIEAQQHL | 0.204545455 |
| B44:02 | RL11 | RDYVDRFYKTL | 0.106918239 | A02:01 | RI9 | RILQQLLFI | 0.201117318 |
| A03:01 | RK10 | RIRTWKSLVK | 0.106382979 | A68:01 | DR11 | DTWAGVEAIIR | 0.2 |
| B14:02 | CC9 | CRAPRKKGC | 0.102564103 | A30:02 | KQY9 | KYCWNLLQY | 0.2 |
| B35:01 | HY9 | HPDIVIYQY | 0.1 | A02:01 | SAV10 | SLLNATAIAV | 0.2 |
| B40:01 | IL8_V4Y | IETYPVKL | 0.1 | A03:01 | RR11 | RLRDLLLIVTR | 0.194174757 |
| C01:02 | VL8 | VIPMFSAL | 0.1 | A01:01 | GY9 | GSEELRSLY | 0.192771084 |
| B40:02 | KA9 | KETINEEAA | 0.098765432 | B40:01 | SL9 | SEGATPQDL | 0.19047619 |
| B40:01 | QL10 | QELKNSAVSL | 0.096385542 | B07:02 | HA9 | HPVHAGPIA | 0.188679245 |
| B07:02 | TM9 | TPQVPLRPM | 0.095238095 | B44:03 | RL11 | RDYVDRFYKTL | 0.186046512 |
| A01:01 | IY9 | ISERILSTY | 0.092592593 | B08:01 | WM8 | WPTVRERM | 0.186046512 |
| A01:01 | YT9 | YFPDWQNYT | 0.092592593 | C12:03 | CC8 | CCFHCQVC | 0.181818182 |
| B15:01 | IY10 | IKLEPVHGVY | 0.092105263 | A68:01 | DWL9 | DTVLEEWNL | 0.181818182 |
| A24:02 | LY10 | LFCASDAKAY | 0.091836735 | B08:01 | GL8 | GGKKKYKL | 0.181818182 |
| B08:01 | WM8_E6A | WPTVRARM | 0.090909091 | B51:01 | EI9 | EKEGKISKI | 0.177777778 |
| A02:01 | YI9_Y9T | YTAFTIPST | 0.090909091 | B51:01 | LI9 | LPCRIKQII | 0.177777778 |
| A30:01 | IY9 | IVNRNRQGY | 0.088235294 | B57:03 | KY10 | KAVRLIKFLY | 0.176470588 |
| A30:01 | KY9 | KQNPDIVIY | 0.088235294 | C03:03 | RL9 | RYLKDQQLL | 0.173913043 |
| A30:01 | KQY9 | KYCWNLLQY | 0.088235294 | A30:01 | KYY9 | KIQNFRVYY | 0.171428571 |
| A68:01 | QV9 | QVSQNYPIV | 0.085714286 | B07:02 | FL9 | FPRIWLHGL | 0.169811321 |
| A03:01 | AK11 | ALVEICTEMEK | 0.085106383 | A02:01 | YI9 | YTAFTIPSI | 0.163841808 |
| B08:01 | DL9 | DCKTILKAL | 0.084745763 | B57:03 | YY9 | YTPGPGIRY | 0.156862745 |
| C04:01 | QW9 | QASQEVKNW | 0.084745763 | B35:01 | DL9 | DPNPQEVVL | 0.155555556 |
| A02:01 | YL9 | YVDRFYKTL | 0.083850932 | B07:02 | SV9 | SPRTLNAWV | 0.150943396 |
| A11:01 | IK10 | IYQEPFKNLK | 0.081081081 | B07:02 | TM9 | TPQVPLRPM | 0.150943396 |
| B40:01 | LS9 | LEKHGAITS | 0.081081081 | B44:03 | AY10 | AENLWVTVYY | 0.148148148 |
| C03:04 | YL9 | YVDRFFKTL | 0.081081081 | A03:01 | ER10 | ERILSTYLGR | 0.14 |
| A02:01 | IV9 | ILKEPVHGV | 0.080745342 | A02:01 | VV9 | VLAEAMSQV | 0.139664804 |
| B57:01 | KI8 | KAFSPEVI | 0.08 | A03:01 | AK11 | ALVEICTEMEK | 0.138613861 |
| B07:02 | SV9 | SPRTLNAWV | 0.079365079 | B57:03 | KI8 | KAFSPEVI | 0.137254902 |
| B18:01 | LY10 | LADQLIHLHY | 0.079365079 | B57:03 | LL9 | LTFGWCFKL | 0.137254902 |
| B51:01 | RL9 | RAIEAQQHL | 0.078125 | B57:03 | QW9 | QASQEVKNW | 0.137254902 |
| A30:01 | KY11 | KQNPDIVIYQY | 0.076923077 | B44:03 | AY9 | AENLWVTVY | 0.136363636 |
| B08:01 | GK9 | GGKKKYKLK | 0.076271186 | B08:01 | DL9 | DCKTILKAL | 0.136363636 |
| A02:01 | RA9 | RIRQGLERA | 0.07165109 | B08:01 | EV9 | ELRSLYNTV | 0.136363636 |
| A03:01 | HK9 | HMYISKKAK | 0.071428571 | B08:01 | GK9 | GGKKKYKLK | 0.136363636 |
| C08:02 | HL9 | HLVWASREL | 0.071428571 | A03:01 | TK10 | TVYYGVPVWK | 0.13592233 |
| A02:01 | SAV10 | SLLNATAIAV | 0.071428571 | A02:01 | RI10 | RGPGRAFVTI | 0.134831461 |
| A03:01 | AK9 | AIFQSSMTK | 0.070921986 | A26:01 | ER11 | ETFYVDGAANR | 0.133333333 |
| A23:01 | RL9 | RYLKDQQLL | 0.068965517 | A30:02 | HY9 | HIGPGRAFY | 0.133333333 |
| A02:01 | VV9 | VLAEAMSQV | 0.068322981 | B57:01 | KF9 | KTAVQMAVF | 0.13253012 |
| A01:01 | WH10 | WRFDSRLAFH | 0.067901235 | B44:02 | LY9 | LYNTVATLY | 0.122807018 |
| B15:03 | WF9 | WRFDSRLAF | 0.065789474 | A02:01 | YL9 | YVDRFYKTL | 0.121621622 |
| A03:01 | RK11 | RMRGAHTNDVK | 0.063829787 | B44:02 | RL11 | RDYVDRFYKTL | 0.120689655 |
| A24:02 | YL8 | YLKDQQLL | 0.06122449 | A01:01 | IY9 | ISERILSTY | 0.120481928 |
| B57:01 | KF9 | KTAVQMAVF | 0.06 | A01:01 | RY9 | RRGWEVLKY | 0.120481928 |
| B08:01 | GL8 | GGKKKYKL | 0.06 | A03:01 | GK9 | GIPHPAGLK | 0.12 |
| A68:01 | IL9 | IVTRIVELL | 0.057142857 | B51:01 | II10 | IPLGDAKLII | 0.119047619 |
| A03:01 | RR11 | RLRDLLLIVTR | 0.056737589 | A11:01 | GK8 | GVGGPGHK | 0.117647059 |
| B07:02 | HA9 | HPVHAGPIA | 0.055555556 | B57:03 | KF9 | KTAVQMAVF | 0.117647059 |
| B07:02 | TL9 | TPQDLNTML | 0.055555556 | B57:03 | KV8 | KAIGTVLV | 0.117647059 |
| A11:01 | AK10 | AVFIHNFKRK | 0.054054054 | B15:01 | RA9 | RMRRAEPAA | 0.117647059 |
| A11:01 | PK8 | PLRPMTYK | 0.054054054 | B44:03 | EV9 | EEKAFSPEV | 0.11627907 |

| | | | |
|---|---|---|---|
| A11:01 | QKK9 | QIIEQLIKK | 0.054054054 |
| A11:01 | SK9 | SVITQACPK | 0.054054054 |
| A02:01 | PL10 | PLTFGWCYKL | 0.052795031 |
| A68:01 | EV10 | ETYGDTWTGV | 0.052631579 |
| B15:10 | YL9 | YVDRFFKTL | 0.052631579 |
| B15:01 | RA9 | RMRRAEPAA | 0.050632911 |
| A02:01 | LV10 | LTFGWCFKLV | 0.049689441 |
| A02:01 | LI9 | LVGPTPVNI | 0.049689441 |
| A03:01 | ER10 | ERILSTYLGR | 0.04964539 |
| A03:01 | GK9 | GIPHPAGLK | 0.04964539 |
| B18:01 | FK10 | FRDYVDRFYK | 0.047619048 |
| B15:10 | HL9 | HQAISPRTL | 0.046875 |
| A02:01 | VL9 | VIYQYMDDL | 0.046583851 |
| A02:01 | AM9 | ALVEICTEM | 0.043478261 |
| A03:01 | KK11 | KTKPPLPSVKK | 0.042857143 |
| A68:01 | IV9 | ITLWQRPLV | 0.042857143 |
| B08:01 | GL9 | GPKVKQWPL | 0.042372881 |
| B08:01 | YL8 | YLKDQQLL | 0.042372881 |
| A11:01 | GK8 | GVGGPGHK | 0.040540541 |
| B57:01 | AW9 | AVRHFPRIW | 0.04 |
| B15:01 | TY11 | TQGYFPDWQNY | 0.039473684 |
| B15:03 | RY9 | RKAKIIRDY | 0.039473684 |
| B40:01 | SL9 | SEGATPQDL | 0.036144578 |
| B35:01 | VL11 | VPVWKEATTTL | 0.033898305 |
| B08:01 | EV9 | ELRSLYNTV | 0.033898305 |
| A26:01 | ER11 | ETFYVDGAANR | 0.03125 |
| B15:10 | FY10 | FKRKGGIGGY | 0.03125 |
| B15:10 | GL9 | GHQAAMQML | 0.03125 |
| B44:02 | EV9 | EEKAFSPEV | 0.030864198 |
| A30:01 | HY9 | HIGPGRAFY | 0.029411765 |
| A30:01 | RV10 | RMRGAHTNDV | 0.029411765 |
| A68:01 | DL9 | DTVLEEMNL | 0.028571429 |
| A03:01 | TK10 | TVYYGVPVWK | 0.028368794 |
| B15:01 | LY12 | LVGKLNWASQIY | 0.026315789 |
| B15:03 | VI10 | VTDSQYALGI | 0.026315789 |
| B35:01 | TW9 | TAVPWNASW | 0.025423729 |
| B35:01 | WF9 | WASRELERF | 0.025423729 |
| B08:01 | RL9 | RVKEKYQHL | 0.025423729 |
| C03:03 | YL9 | YVDRFFKTL | 0.024390244 |
| C07:01 | KY11_K1R | RRQEILDLWVY | 0.023255814 |
| B57:01 | FF9 | FSPEVIPMF | 0.023255814 |
| B57:01 | KV8 | KAIGTVLV | 0.023255814 |
| B57:01 | LL9 | LTFGWCFKL | 0.023255814 |
| A02:01 | VL10 | VLEWRFDSRL | 0.02173913 |
| A03:01 | AK10 | AVFIHNFKRK | 0.021276596 |
| A03:01 | KK10 | KLVDFRELNK | 0.021276596 |
| B57:01 | YT9 | YFPDWQNYT | 0.02 |
| A02:01 | KL9 | KLTPLCVTL | 0.01863354 |
| A02:01 | RI10 | RGPGRAFVTI | 0.01863354 |
| B51:01 | II10 | IPLGDAKLII | 0.018181818 |
| B07:02 | FR10 | FPVTPQVPLR | 0.017857143 |
| B07:02 | FL9 | FPVTPQVPL | 0.015873016 |
| B15:03 | IY9 | IQQEFGIPY | 0.015625 |
| B15:03 | VF9 | VKVIEEKAF | 0.015625 |
| A11:01 | TI9 | TLYCVHQRI | 0.013513514 |
| B35:01 | NY9 | NSSKVSQNY | 0.008474576 |
| B35:01 | NIY9 | NPVPVGNIY | 0 |
| A33:01 | ER8 | EVAQRAYR | 0 |
| A33:01 | VR10 | VFAVLSIVNR | 0 |
| A30:01 | LF11 | LVWASRELERF | 0 |
| B15:10 | WI8 | WHLGHVSI | 0 |
| B15:16 | SF9 | SFNCGGEFF | 0 |
| A23:01 | LY9 | LRSLFLFSY | 0 |
| B55:01 | VT10 | VPVWKEATTT | 0 |

| | | | |
|---|---|---|---|
| B08:01 | YL8 | YLKDQQLL | 0.11627907 |
| A30:01 | KIY9 | KLNWASQIY | 0.114285714 |
| A30:01 | KY11 | KEKGGLEGLIY | 0.114285714 |
| A03:01 | KK11 | KTKPPLPSVKK | 0.111111111 |
| A03:01 | HK9 | HMYISKKAK | 0.11 |
| A03:01 | QR9 | QIYPGIKVR | 0.108910891 |
| B35:01 | VL11 | VPVWKEATTTL | 0.108695652 |
| B35:01 | TW9 | TAVPWNASW | 0.106382979 |
| B57:01 | YY9 | YTPGPGIRY | 0.1 |
| A02:01 | KL9 | KLTPLCVTL | 0.095505618 |
| A02:01 | VL9 | VIYQYMDDL | 0.08988764 |
| A30:02 | IY9 | IVNRNRQGY | 0.088888889 |
| A30:01 | IY9 | IVNRNRQGY | 0.088235294 |
| A02:01 | LV10 | LTFGWCFKLV | 0.083333333 |
| B57:03 | IW9 | IVLPEKDSW | 0.078431373 |
| C08:02 | SL9 | SAEPVPLQL | 0.076923077 |
| B57:01 | FF9 | FSPEVIPMF | 0.075 |
| A11:01 | SK9 | SVITQACPK | 0.074074074 |
| A68:02 | IL9 | IVTRIVELL | 0.071428571 |
| A03:01 | KK10 | KLVDFRELNK | 0.07 |
| A68:02 | IV9 | ITLWQRPLV | 0.068965517 |
| A68:02 | QV9 | QVSQNYPIV | 0.068965517 |
| A02:01 | VL10 | VLEWRFDSRL | 0.06741573 |
| A02:01 | LI9 | LVGPTPVNI | 0.067039106 |
| A68:02 | DL9 | DTVLEEMNL | 0.066666667 |
| B14:02 | SL9 | SAEPVPLQL | 0.066666667 |
| C05:01 | SL9 | SAEPVPLQL | 0.066666667 |
| B18:01 | FK10 | FRDYVDRFYK | 0.0625 |
| A11:01 | GK9 | GIPHPAGLK | 0.0625 |
| A02:01 | AM9 | ALVEICTEM | 0.061797753 |
| A01:01 | WH10 | WRFDSRLAFH | 0.060240964 |
| B57:03 | IF9 | ISKKAKGWF | 0.058823529 |
| A30:01 | KQY9 | KYCWNLLQY | 0.057142857 |
| A30:01 | RV10 | RMRGAHTNDV | 0.057142857 |
| B07:02 | FL9 | FPVTPQVPL | 0.056603774 |
| A11:01 | TI9 | TLYCVHQRI | 0.055555556 |
| B44:02 | AY9 | AENLWVTVY | 0.051724138 |
| B57:01 | KV8 | KAIGTVLV | 0.05 |
| B35:01 | PY9 | PPIPVGDIY | 0.043478261 |
| A68:02 | DWL9 | DTVLEEWNL | 0.04 |
| A68:01 | QV9 | QVSQNYPIV | 0.04 |
| B07:02 | FR10 | FPVTPQVPLR | 0.038461538 |
| B57:01 | LL9 | LTFGWCFKL | 0.037974684 |
| B57:01 | KI8 | KAFSPEVI | 0.036585366 |
| B44:02 | EV9 | EEKAFSPEV | 0.034482759 |
| A30:01 | HY9 | HIGPGRAFY | 0.028571429 |
| A30:01 | KY9 | KQNPDIVIY | 0.028571429 |
| B53:01 | TL9 | TPYDINQML | 0.026315789 |
| B08:01 | RL9 | RVKEKYQHL | 0.022727273 |
| B35:01 | NY9 | NSSKVSQNY | 0.02173913 |
| A11:01 | PK8 | PLRPMTYK | 0.018181818 |
| B44:02 | AY10 | AENLWVTVYY | 0 |
| A68:01 | DL9 | DTVLEEMNL | 0 |
| A11:01 | IK9 | IIATDIQTK | 0 |
| A68:01 | IL9 | IVTRIVELL | 0 |
| A68:01 | IV9 | ITLWQRPLV | 0 |
| A23:01 | LY9 | LRSLFLFSY | 0 |

### Supplementary Table S2

| HLA | Peptide Sequence | Protein | Amplitude | Source |
|---|---|---|---|---|
| HLA-A*01:01 | ISDYDYYRY | orf1ab | 1712222.222 | Prachar et al; Ahmed et al |
| HLA-A*11:01 | KTFPPTEPK | N | 1217377.049 | Prachar et al; Ahmed et al |
| HLA-A*03:01 | KTFPPTEPKK | N | 1071000 | Ahmed et al |
| HLA-A*24:02 | NYMPYFFTL | orf1ab | 1050428.571 | Prachar et al |
| HLA-A*01:01 | LTDEMIAQY | S | 953846.1538 | Prachar et al |
| HLA-A*01:01 | DTDFVNEFY | orf1ab | 911428.5714 | Prachar et al |
| HLA-A*03:01 | KTFPPTEPK | N | 894698.7952 | Prachar et al; Ahmed et al |
| HLA-A*01:01 | GTDLEGNFY | orf1ab | 827560.9756 | Prachar et al |
| HLA-A*11:01 | STFNVPMEK | orf1ab | 805555.5556 | Prachar et al |
| HLA-A*01:01 | CTDDNALAY | orf1ab | 787500 | Prachar et al |
| HLA-A*11:01 | TTIKPVTYK | orf1ab | 688235.2941 | Prachar et al |
| HLA-A*24:02 | YYTSNPTTF | orf1ab | 625781.25 | Prachar et al |
| HLA-A*02:03 | FLLPSLATV | orf1ab | 513518.5185 | Ahmed et al |
| HLA-A*11:01 | KTFPPTEPKK | N | 507315.7895 | Ahmed et al |
| HLA-A*01:01 | FTSDYYQLY | orf3a | 418524.5902 | Prachar et al |
| HLA-B*54:01 | FPRGQGVPI | N | 366701.5707 | Ahmed et al |
| HLA-B*15:01 | VVYRGTTTY | orf1ab | 352592.5926 | Ahmed et al |
| HLA-A*11:01 | TSFGPLVRK | orf1ab | 342592.5926 | Prachar et al |
| HLA-B*27:05 | VRFPNITNL | S | 338437.5 | Ahmed et al |
| HLA-A*01:01 | PTDNYITTY | orf1ab | 337066.6667 | Prachar et al |
| HLA-A*03:01 | STFNVPMEK | orf1ab | 329545.4545 | Prachar et al |
| HLA-A*24:02 | VYMPASWVM | orf1ab | 324310.3448 | Prachar et al |
| HLA-B*07:02 | FPRGQGVPI | N | 322764.977 | Ahmed et al |
| HLA-A*11:01 | ASMPTTIAK | orf1ab | 301764.7059 | Prachar et al |
| HLA-B*15:01 | RLYYDSMSY | orf1ab | 293333.3333 | Ahmed et al |
| HLA-A*11:01 | KSAGFPFNK | orf1ab | 281538.4615 | Prachar et al |
| HLA-A*02:06 | FLLPSLATV | orf1ab | 274554.4554 | Ahmed et al |
| HLA-C*07:01 | KRVDWTIEY | orf1ab | 260653.0612 | Prachar et al |
| HLA-A*03:01 | TTIKPVTYK | orf1ab | 254347.8261 | Prachar et al |
| HLA-A*02:01 | FLLPSLATV | orf1ab | 247589.2857 | Prachar et al; Ahmed et al |
| HLA-A*24:02 | YYHTTDPSF | orf1ab | 247011.9522 | Prachar et al |
| HLA-B*40:01 | GEYSHVVAF | orf1ab | 239811.3208 | Prachar et al |
| HLA-A*02:06 | FVDGVPFVV | orf1ab | 238333.3333 | Ahmed et al |
| HLA-A*03:01 | KLFDRYFKY | orf1ab | 230769.2308 | Prachar et al |
| HLA-A*02:01 | VLWAHGFEL | orf1ab | 214698.7952 | Ahmed et al |
| HLA-A*03:01 | RLYYDSMSY | orf1ab | 209523.8095 | Ahmed et al |
| HLA-B*40:01 | SEVGPEHSL | orf1ab | 198857.1429 | Prachar et al |
| HLA-A*11:01 | HVVGPNVNK | orf1ab | 193129.771 | Prachar et al |
| HLA-A*02:02 | FLLPSLATV | orf1ab | 191241.3793 | Ahmed et al |
| HLA-A*02:06 | LLLDDFVEI | orf1ab | 190000 | Ahmed et al |
| HLA-C*07:02 | KRVDWTIEY | orf1ab | 189495.549 | Prachar et al |
| HLA-A*02:01 | LLLDDFVEI | orf1ab | 169586.7769 | Ahmed et al |
| HLA-A*02:01 | YLQPRTFLL | S | 154000 | Prachar et al |
| HLA-B*40:01 | GETLPTEVL | orf1ab | 153141.3613 | Prachar et al |
| HLA-A*01:01 | VTPSGTWLTY | N | 151911.2207 | Ahmed et al |
| HLA-A*02:01 | ALWEIQQVV | orf1ab | 148780.4878 | Ahmed et al |

| | | | | |
|---|---|---|---|---|
| HLA-B*40:01 | QEYADVFHL | orf1ab | 148625 | Prachar et al |
| HLA-A*02:02 | TMADLVYAL | orf1ab | 139830.5085 | Ahmed et al |
| HLA-A*02:01 | FLAHIQWMV | orf1ab | 136012.4611 | Prachar et al |
| HLA-A*02:01 | FLLNKEMYL | orf1ab | 125517.2414 | Prachar et al; Ahmed et al |
| HLA-A*03:01 | QIYKTPPIK | S | 123378.3784 | Prachar et al |
| HLA-A*02:01 | YLDAYNMMI | orf1ab | 120215.0538 | Prachar et al; Ahmed et al |
| HLA-A*03:01 | KSAGFPFNK | orf1ab | 119024.3902 | Prachar et al |
| HLA-A*02:02 | FLLNKEMYL | orf1ab | 117419.3548 | Ahmed et al |
| HLA-A*11:01 | AVAKHDFFK | orf1ab | 110000 | Prachar et al |
| HLA-B*07:02 | IPRRNVATL | orf1ab | 107663.5514 | Ahmed et al |
| HLA-A*11:01 | AVLQSGFRK | orf1ab | 106250 | Prachar et al; Ahmed et al |
| HLA-A*11:01 | GVAMPNLYK | orf1ab | 99836.06557 | Prachar et al; Ahmed et al |
| HLA-A*02:01 | FVDGVPFVV | orf1ab | 99535.96288 | Ahmed et al |
| HLA-B*40:01 | MEVTPSGTWL | N | 97994.29658 | Ahmed et al |
| HLA-A*24:02 | FYGGWHNML | orf1ab | 96814.44992 | Prachar et al |
| HLA-C*06:02 | FRYMNSQGL | orf1ab | 96470.58824 | Ahmed et al |
| HLA-A*11:01 | VTNNTFTLK | orf1ab | 96325.3012 | Prachar et al |
| HLA-B*51:01 | FPRGQGVPI | N | 92401.05541 | Ahmed et al |
| HLA-A*11:01 | VTYVPAQEK | S | 91022.22222 | Prachar et al |
| HLA-A*03:01 | VTYVPAQEK | S | 90220.26432 | Prachar et al |
| HLA-A*03:01 | TSFGPLVRK | orf1ab | 86854.46009 | Prachar et al |
| HLA-A*30:02 | VTPSGTWLTY | N | 82490.79344 | Ahmed et al |
| HLA-A*24:02 | NYNYLYRLF | S | 80769.23077 | Prachar et al |
| HLA-A*02:06 | YLDAYNMMI | orf1ab | 80431.65468 | Ahmed et al |
| HLA-A*02:06 | VLWAHGFEL | orf1ab | 79910.3139 | Ahmed et al |
| HLA-A*01:01 | LTNDNTSRY | orf1ab | 77924.5283 | Prachar et al |
| HLA-A*02:02 | VLWAHGFEL | orf1ab | 77478.26087 | Ahmed et al |
| HLA-A*03:01 | VVYRGTTTY | orf1ab | 75957.44681 | Ahmed et al |
| HLA-A*02:06 | FLLNKEMYL | orf1ab | 74285.71429 | Ahmed et al |
| HLA-A*29:02 | VTPSGTWLTY | N | 72236.88068 | Ahmed et al |
| HLA-A*24:02 | VYSTGSNVF | S | 70904.25532 | Prachar et al |
| HLA-C*04:01 | GFDYVYNPF | orf1ab | 69349.31507 | Prachar et al |
| HLA-A*02:02 | YLDAYNMMI | orf1ab | 69012.34568 | Ahmed et al |
| HLA-A*02:02 | LLLDDFVEI | orf1ab | 68859.0604 | Ahmed et al |
| HLA-B*15:01 | YQKVGMQKY | orf1ab | 68055.55556 | Ahmed et al |
| HLA-A*03:01 | GVAMPNLYK | orf1ab | 67666.66667 | Prachar et al; Ahmed et al |
| HLA-A*02:06 | ALWEIQQVV | orf1ab | 67477.87611 | Ahmed et al |
| HLA-A*03:01 | IINNTVYTK | orf1ab | 63377.77778 | Prachar et al |
| HLA-B*27:05 | GRLQSLQTY | S | 63267.97386 | Ahmed et al |
| HLA-B*40:01 | HEETIYNLL | orf1ab | 61746.36175 | Prachar et al |
| HLA-A*02:02 | FLNRFTTTL | orf1ab | 61327.43363 | Ahmed et al |
| HLA-A*03:01 | ASMPTTIAK | orf1ab | 59651.16279 | Prachar et al |
| HLA-A*02:03 | FLLNKEMYL | orf1ab | 56875 | Ahmed et al |
| HLA-A*26:01 | VTPSGTWLTY | N | 56449.02635 | Ahmed et al |
| HLA-A*02:01 | TMADLVYAL | orf1ab | 56122.44898 | Ahmed et al |
| HLA-A*24:02 | YYKKDNSYF | orf1ab | 55706.80628 | Prachar et al |
| HLA-A*11:01 | AIDAYPLTK | orf1ab | 55354.33071 | Prachar et al |
| HLA-A*24:02 | TYKPNTWCI | orf1ab | 54453.26279 | Prachar et al |
| HLA-A*11:01 | IINNTVYTK | orf1ab | 53208.95522 | Prachar et al |

| | | | | |
|---|---|---|---|---|
| HLA-A*30:02 | ISDYDYYRY | orf1ab | 53092.16193 | Ahmed et al |
| HLA-A*02:02 | ALWEIQQVV | orf1ab | 52495.69707 | Ahmed et al |
| HLA-A*03:01 | VTNNTFTLK | orf1ab | 51747.57282 | Prachar et al |
| HLA-A*02:03 | FLNRFTTTL | orf1ab | 51716.41791 | Ahmed et al |
| HLA-A*03:01 | GVYFASTEK | S | 49354.83871 | Prachar et al |
| HLA-B*58:01 | KQFDTYNLW | orf1ab | 49034.58213 | Ahmed et al |
| HLA-A*03:01 | TLKGGAPTK | orf1ab | 48829.78723 | Prachar et al |
| HLA-B*07:02 | SPRWYFYYL | N | 48653.84615 | Ahmed et al |
| HLA-B*40:01 | HEGKTFYVL | orf1ab | 46976.74419 | Prachar et al |
| HLA-C*14:02 | VRFPNITNL | S | 45928.75318 | Ahmed et al |
| HLA-A*02:06 | TMADLVYAL | orf1ab | 44836.95652 | Ahmed et al |
| HLA-A*31:01 | KTFPPTEPK | N | 44600.6006 | Ahmed et al |
| HLA-B*40:01 | YEGNSPFHPL | orf7a | 44216.07857 | Ahmed et al |
| HLA-A*03:01 | HLYLQYIRK | orf1ab | 43893.12977 | Prachar et al |
| HLA-A*02:01 | LLLDRLNQL | N | 43120.56738 | Ahmed et al |
| HLA-A*03:01 | ALAYYNTTK | orf1ab | 42612.61261 | Prachar et al |
| HLA-A*02:03 | ALWEIQQVV | orf1ab | 41666.66667 | Ahmed et al |
| HLA-A*03:01 | RIAGHHLGR | M | 41198.97959 | Prachar et al |
| HLA-A*01:01 | VVDKYFDCY | orf1ab | 40993.15068 | Prachar et al; Ahmed et al |
| HLA-A*11:01 | HLMGWDYPK | orf1ab | 40745.68544 | Prachar et al |
| HLA-A*29:02 | ISDYDYYRY | orf1ab | 40499.34297 | Ahmed et al |
| HLA-A*03:01 | VLSGHNLAK | orf1ab | 40000 | Prachar et al |
| HLA-A*30:02 | GTTLPKGFY | N | 39809.34223 | Ahmed et al |
| HLA-A*29:02 | TPSGTWLTY | N | 39157.09558 | Ahmed et al |
| HLA-A*02:02 | FVDGVPFVV | orf1ab | 37764.08451 | Ahmed et al |
| HLA-A*11:01 | GVYFASTEK | S | 35581.39535 | Prachar et al |
| HLA-A*11:01 | KTIQPRVEK | orf1ab | 35573.77049 | Prachar et al |
| HLA-A*02:01 | VLNDILSRL | S | 34887.64045 | Ahmed et al |
| HLA-A*02:03 | TMADLVYAL | orf1ab | 34810.12658 | Ahmed et al |
| HLA-A*02:01 | ALSKGVHFV | orf3a | 34705.88235 | Prachar et al |
| HLA-A*68:02 | FVDGVPFVV | orf1ab | 34485.53055 | Ahmed et al |
| HLA-A*30:02 | ATSRTLSYY | M | 34156.62651 | Ahmed et al |
| HLA-A*02:03 | LLLDDFVEI | orf1ab | 33917.35537 | Ahmed et al |
| HLA-B*58:01 | TTLPVNVAF | orf1ab | 33585.78053 | Ahmed et al |
| HLA-A*02:02 | HLVDFQVTI | orf6 | 33333.33333 | Ahmed et al |
| HLA-A*02:03 | FLPRVFSAV | orf1ab | 32927.46114 | Ahmed et al |
| HLA-B*40:01 | YELQTPFEI | orf1ab | 32640 | Prachar et al |
| HLA-B*40:01 | QELYSPIFL | orf7a | 32178.42324 | Prachar et al |
| HLA-A*02:01 | TLVPQEHYV | orf1ab | 31550.3876 | Ahmed et al |
| HLA-B*18:01 | YEQYIKWPWY | S | 31476.23019 | Ahmed et al |
| HLA-A*24:02 | IYNDKVAGF | orf1ab | 31232.87671 | Prachar et al |
| HLA-B*40:01 | AEWFLAYIL | orf1ab | 30457.22714 | Prachar et al |
| HLA-B*35:01 | VVYRGTTTY | orf1ab | 30222.22222 | Ahmed et al |
| HLA-B*40:01 | RELHLSWEV | orf1ab | 29360.04162 | Prachar et al |
| HLA-A*01:01 | LIDLQELGKY | S | 29204.66596 | Ahmed et al |
| HLA-A*11:01 | VVYRGTTTY | orf1ab | 29024.39024 | Ahmed et al |
| HLA-A*03:01 | AVLQSGFRK | orf1ab | 28419.45289 | Ahmed et al |
| HLA-A*24:02 | FYLTNDVSF | orf1ab | 28249.25816 | Prachar et al |
| HLA-B*40:01 | SEFDRDAAM | orf1ab | 27952.21843 | Prachar et al |

| | | | | |
|---|---|---|---|---|
| HLA-B*07:02 | HPLADNKFAL | orf7a | 27222.22222 | Ahmed et al |
| HLA-C*07:01 | VYMPASWVM | orf1ab | 26833.09558 | Prachar et al |
| HLA-A*11:01 | AGFSLWVYK | orf1ab | 26812.5 | Prachar et al |
| HLA-A*02:03 | KLSYGIATV | orf1ab | 25517.24138 | Ahmed et al |
| HLA-A*24:02 | SYATHSDKF | orf1ab | 25158.06988 | Prachar et al |
| HLA-A*02:01 | ILGTVSWNL | orf1ab | 25039.44316 | Ahmed et al |
| HLA-A*23:01 | LWLLWPVTL | M | 24662.73187 | Ahmed et al |
| HLA-A*68:01 | KTFPPTEPK | N | 24459.81555 | Ahmed et al |
| HLA-B*40:01 | GEVITFDNL | orf1ab | 24285.71429 | Prachar et al |
| HLA-B*44:03 | YEQYIKWPWY | S | 23844.57432 | Ahmed et al |
| HLA-B*40:02 | YEGNSPFHPL | orf7a | 23515.18669 | Ahmed et al |
| HLA-A*24:02 | TYACWHHSI | orf1ab | 23066.92913 | Prachar et al |
| HLA-B*44:02 | YEQYIKWPWY | S | 22942.24924 | Ahmed et al |
| HLA-A*02:01 | YLYALVYFL | orf3a | 22906.50407 | Prachar et al |
| HLA-A*02:01 | HLVDFQVTI | orf6 | 22500 | Ahmed et al |
| HLA-B*40:01 | TEHSWNADL | orf1ab | 22258.45903 | Prachar et al |
| HLA-A*02:06 | LVLSVNPYV | orf1ab | 22058.82353 | Ahmed et al |
| HLA-A*02:06 | YTMADLVYA | orf1ab | 21982.50729 | Ahmed et al |
| HLA-A*24:02 | LWLLWPVTL | M | 21731.05498 | Ahmed et al |
| HLA-B*40:01 | AEYHNESGL | orf1ab | 20615.18325 | Prachar et al |
| HLA-A*02:03 | VLWAHGFEL | orf1ab | 20490.60943 | Ahmed et al |
| HLA-A*02:01 | LLDDFVEII | orf1ab | 20276.72956 | Ahmed et al |
| HLA-B*40:01 | TEVVGDIIL | orf1ab | 19889.38053 | Prachar et al |
| HLA-B*07:02 | FPPTSFGPL | orf1ab | 19859.15493 | Ahmed et al |
| HLA-A*03:01 | AVAKHDFFK | orf1ab | 19446.42857 | Prachar et al |
| HLA-B*40:01 | FELEDFIPM | orf1ab | 19400.35273 | Prachar et al |
| HLA-A*02:01 | NLIDSYFVV | orf1ab | 19288.99083 | Prachar et al |
| HLA-A*01:01 | LSPRWYFYY | N | 18750 | Ahmed et al |
| HLA-A*31:01 | SVSPKLFIR | orf7a | 18576.51246 | Ahmed et al |
| HLA-A*02:02 | FLPRVFSAV | orf1ab | 18028.36879 | Ahmed et al |
| HLA-A*02:01 | FLNRFTTTL | orf1ab | 17953.36788 | Prachar et al; Ahmed et al |
| HLA-A*02:06 | GVYDYLVST | orf1ab | 17883.75559 | Ahmed et al |
| HLA-A*02:03 | YLDAYNMMI | orf1ab | 17253.08642 | Ahmed et al |
| HLA-A*02:03 | YLNTLTLAV | orf1ab | 16842.10526 | Ahmed et al |
| HLA-A*31:01 | KTFPPTEPKK | N | 16708.26833 | Ahmed et al |
| HLA-A*24:02 | YFIASFRLF | M | 16649.48454 | Prachar et al |
| HLA-A*03:01 | KLFAAETLK | orf1ab | 16551.72414 | Prachar et al; Ahmed et al |
| HLA-A*02:03 | HLVDFQVTI | orf6 | 16527.54591 | Ahmed et al |
| HLA-B*08:01 | FPRGQGVPI | N | 16356.8426 | Ahmed et al |
| HLA-A*68:02 | YTMADLVYA | orf1ab | 16042.55319 | Ahmed et al |
| HLA-A*24:02 | FFASFYYVW | orf1ab | 16003.18471 | Prachar et al |
| HLA-B*35:01 | FPRGQGVPI | N | 15936.29124 | Ahmed et al |
| HLA-A*03:01 | RLFRKSNLK | S | 15810.81081 | Prachar et al |
| HLA-B*40:01 | AELAKNVSL | orf1ab | 15555.55556 | Prachar et al |
| HLA-A*11:01 | GVYYHKNNK | S | 15455.19713 | Prachar et al |
| HLA-B*58:01 | VVYRGTTTY | orf1ab | 15404.53074 | Ahmed et al |
| HLA-A*24:02 | AYANSVFNI | orf1ab | 14970.05988 | Prachar et al |
| HLA-A*02:01 | YLNSTNVTI | orf1ab | 14911.2426 | Prachar et al |
| HLA-A*29:02 | LSPRWYFYY | N | 14877.62988 | Ahmed et al |

| | | | | |
|---|---|---|---|---|
| HLA-A*02:01 | NLWNTFTRL | orf1ab | 14293.40511 | Ahmed et al |
| HLA-A*02:06 | IQPGQTFSV | orf1ab | 14136.80782 | Ahmed et al |
| HLA-A*24:02 | VYFLQSINF | orf3a | 14000 | Prachar et al |
| HLA-A*01:01 | VTDVTQLYL | orf1ab | 13877.55102 | Prachar et al |
| HLA-A*03:01 | RQFHQKLLK | orf1ab | 13562.38698 | Prachar et al |
| HLA-A*02:06 | HLVDFQVTI | orf6 | 13487.73842 | Ahmed et al |
| HLA-A*02:02 | LLDDFVEII | orf1ab | 13137.73431 | Ahmed et al |
| HLA-A*11:01 | ATSRTLSYYK | M | 12840.17279 | Ahmed et al |
| HLA-A*02:01 | YLNTLTLAV | orf1ab | 12715.23179 | Ahmed et al |
| HLA-A*01:01 | FSAVGNICY | orf1ab | 12633.05322 | Prachar et al |
| HLA-B*15:01 | KQFDTYNLW | orf1ab | 12585.05917 | Ahmed et al |
| HLA-B*40:01 | FEYVSQPFL | S | 12500 | Prachar et al |
| HLA-A*02:03 | FLNGSCGSV | orf1ab | 12340.42553 | Ahmed et al |
| HLA-C*07:01 | HANEYRLYL | orf1ab | 12326.84825 | Prachar et al |
| HLA-B*07:02 | MPASWVMRI | orf1ab | 12213.03949 | Ahmed et al |
| HLA-B*53:01 | FPRGQGVPI | N | 12168.17234 | Ahmed et al |
| HLA-A*11:01 | RLYYDSMSY | orf1ab | 11934.90054 | Ahmed et al |
| HLA-A*02:01 | KLNVGDYFV | orf1ab | 11865.96583 | Ahmed et al |
| HLA-A*02:06 | FLPRVFSAV | orf1ab | 11845.29357 | Ahmed et al |
| HLA-A*24:02 | RYKLEGYAF | orf1ab | 11755.55556 | Prachar et al |
| HLA-A*68:01 | KTFPPTEPKK | N | 11507.87966 | Ahmed et al |
| HLA-A*02:01 | SMWSFNPET | M | 11365.31365 | Ahmed et al |
| HLA-A*02:02 | YLNTLTLAV | orf1ab | 10971.42857 | Ahmed et al |
| HLA-A*02:02 | KLNVGDYFV | orf1ab | 10945.45455 | Ahmed et al |
| HLA-A*11:01 | ALAYYNTTK | orf1ab | 10911.188 | Prachar et al |
| HLA-A*02:03 | FVDGVPFVV | orf1ab | 10679.61165 | Ahmed et al |
| HLA-A*02:01 | FLWLLWPVTL | M | 10564.70351 | Ahmed et al |
| HLA-A*24:02 | SFNPETNIL | M | 10017.71479 | Ahmed et al |
| HLA-A*69:01 | TLVPQEHYV | orf1ab | 9987.730061 | Ahmed et al |
| HLA-A*02:02 | FLGRYMSAL | orf1ab | 9806.2954 | Ahmed et al |
| HLA-B*51:01 | FPPTSFGPL | orf1ab | 9625.217508 | Ahmed et al |
| HLA-A*01:01 | RVDFCGKGY | S | 9577.390386 | Ahmed et al |
| HLA-B*35:01 | MPASWVMRI | orf1ab | 9414.818311 | Ahmed et al |
| HLA-A*24:02 | KQFDTYNLW | orf1ab | 9403.150041 | Ahmed et al |
| HLA-A*02:01 | FLPRVFSAV | orf1ab | 9393.939394 | Ahmed et al |
| HLA-A*01:01 | FLTENLLLY | orf1ab | 9321.382843 | Prachar et al |
| HLA-A*11:01 | ITPVHVMSK | orf1ab | 9176.470588 | Prachar et al |
| HLA-A*01:01 | LTGHMLDMY | orf1ab | 9101.471727 | Prachar et al |
| HLA-B*54:01 | SPRWYFYYL | N | 9088.459811 | Ahmed et al |
| HLA-A*02:01 | LVLSVNPYV | orf1ab | 9068.278805 | Ahmed et al |
| HLA-A*02:01 | WLLWPVTLA | M | 9039.644566 | Ahmed et al |
| HLA-A*02:03 | FLGRYMSAL | orf1ab | 8881.578947 | Ahmed et al |
| HLA-A*24:02 | GYQPYRVVVL | S | 8855.580694 | Ahmed et al |
| HLA-A*11:01 | QTFFKLVNK | orf1ab | 8849.557522 | Prachar et al |
| HLA-A*30:02 | LSPRWYFYY | N | 8754.421425 | Ahmed et al |
| HLA-B*53:01 | FPNITNLCPF | S | 8723.958333 | Ahmed et al |
| HLA-B*35:01 | HPLADNKFAL | orf7a | 8677.520708 | Ahmed et al |
| HLA-B*40:01 | REVLSDREL | orf1ab | 8645.418327 | Prachar et al |
| HLA-A*29:02 | GTTLPKGFY | N | 8617.416426 | Ahmed et al |

| | | | | |
|---|---|---|---|---|
| HLA-B*35:01 | FPNITNLCPF | S | 8555.839331 | Ahmed et al |
| HLA-A*11:01 | SVLNDILSR | S | 8490.566038 | Ahmed et al |
| HLA-A*02:01 | KLWAQCVQL | orf1ab | 8470.319635 | Ahmed et al |
| HLA-A*03:01 | TLKSFTVEK | S | 8461.538462 | Prachar et al |
| HLA-A*30:02 | RVDFCGKGY | S | 8350.990327 | Ahmed et al |
| HLA-A*02:06 | LLDDFVEII | orf1ab | 8309.278351 | Ahmed et al |
| HLA-B*54:01 | APHGVVFLHV | S | 8265.916924 | Ahmed et al |
| HLA-B*40:01 | REQIDGYVM | orf1ab | 8166.666667 | Prachar et al |
| HLA-C*01:02 | FAPSASAFF | N | 8118.959108 | Prachar et al |
| HLA-B*54:01 | FPPTSFGPL | orf1ab | 7998.887653 | Ahmed et al |
| HLA-B*45:01 | AEGSRGGSQA | N | 7948.303716 | Ahmed et al |
| HLA-A*23:01 | SFNPETNIL | M | 7883.364312 | Ahmed et al |
| HLA-B*53:01 | HPLADNKFAL | orf7a | 7842.023748 | Ahmed et al |
| HLA-A*69:01 | VLWAHGFEL | orf1ab | 7822.651449 | Ahmed et al |
| HLA-A*24:02 | YYQLYSTQL | orf3a | 7813.504823 | Prachar et al |
| HLA-A*02:03 | TLIGDCATV | orf1ab | 7774.86911 | Ahmed et al |
| HLA-A*23:01 | GYQPYRVVVL | S | 7618.815896 | Ahmed et al |
| HLA-B*15:01 | TTLPVNVAF | orf1ab | 7603.219034 | Ahmed et al |
| HLA-A*02:01 | RLDKVEAEV | S | 7536.382536 | Ahmed et al |
| HLA-A*68:02 | MVMCGGSLYV | orf1ab | 7314.741036 | Ahmed et al |
| HLA-A*02:02 | KLSYGIATV | orf1ab | 7184.466019 | Ahmed et al |
| HLA-B*44:03 | LEQWNLVIGF | M | 7048.888889 | Ahmed et al |

#### Supplementary Table S3

| HLA | Peptide | Protein | Amplitude | in229E | inOC43 | inHL63 | inHKU1 |
|---|---|---|---|---|---|---|---|
| HLA-A*02:01 | VYLPYPDPSRI | orf1ab | 9433.26433 | FALSE | FALSE | FALSE | TRUE |
| HLA-A*02:01 | YLPYPDPSRIL | orf1ab | 14539.13997 | FALSE | FALSE | TRUE | TRUE |
| HLA-A*02:01 | YLPYPDPSRI | orf1ab | 144083.5267 | TRUE | FALSE | TRUE | TRUE |
| HLA-A*02:01 | YVYIGDPAQL | orf1ab | 9934.695012 | FALSE | TRUE | FALSE | TRUE |
| HLA-A*02:01 | IFVDGVPFV | orf1ab | 10458.07661 | FALSE | TRUE | FALSE | TRUE |
| HLA-A*02:01 | FVDGVPFVV | orf1ab | 99535.96288 | FALSE | TRUE | FALSE | TRUE |
| HLA-A*02:02 | VYLPYPDPSRI | orf1ab | 9849.059278 | FALSE | FALSE | FALSE | TRUE |
| HLA-A*02:02 | YLPYPDPSRIL | orf1ab | 18746.17016 | FALSE | FALSE | TRUE | TRUE |
| HLA-A*02:02 | YLPYPDPSRI | orf1ab | 205970.1493 | TRUE | FALSE | TRUE | TRUE |
| HLA-A*02:02 | YVYIGDPAQL | orf1ab | 14400.80564 | FALSE | TRUE | FALSE | TRUE |
| HLA-A*02:02 | FVDGVPFVV | orf1ab | 37764.08451 | FALSE | TRUE | FALSE | TRUE |
| HLA-A*02:03 | VYLPYPDPSRI | orf1ab | 12416.58105 | FALSE | FALSE | FALSE | TRUE |
| HLA-A*02:03 | YLPYPDPSRIL | orf1ab | 11499.62049 | FALSE | FALSE | TRUE | TRUE |
| HLA-A*02:03 | YLPYPDPSRI | orf1ab | 330758.988 | TRUE | FALSE | TRUE | TRUE |
| HLA-A*02:03 | YVYIGDPAQL | orf1ab | 11588.33063 | FALSE | TRUE | FALSE | TRUE |
| HLA-A*02:03 | FVDGVPFVV | orf1ab | 10679.61165 | FALSE | TRUE | FALSE | TRUE |
| HLA-A*02:06 | YLPYPDPSRIL | orf1ab | 9793.154395 | FALSE | FALSE | TRUE | TRUE |
| HLA-A*02:06 | YLPYPDPSRI | orf1ab | 51016.63586 | TRUE | FALSE | TRUE | TRUE |
| HLA-A*02:06 | YVYIGDPAQL | orf1ab | 17593.50394 | FALSE | TRUE | FALSE | TRUE |
| HLA-A*02:06 | IFVDGVPFV | orf1ab | 16683.76068 | FALSE | TRUE | FALSE | TRUE |
| HLA-A*02:06 | FVDGVPFVV | orf1ab | 238333.3333 | FALSE | TRUE | FALSE | TRUE |
| HLA-A*02:06 | FVDGVPFV | orf1ab | 8002.910149 | FALSE | TRUE | FALSE | TRUE |
| HLA-A*68:02 | YLPYPDPSRI | orf1ab | 10690.76824 | TRUE | FALSE | TRUE | TRUE |
| HLA-A*68:02 | YVYIGDPAQL | orf1ab | 11261.616 | FALSE | TRUE | FALSE | TRUE |
| HLA-A*68:02 | EIVDTVSALV | orf1ab | 10343.47399 | FALSE | TRUE | FALSE | FALSE |
| HLA-A*68:02 | FVDGVPFVV | orf1ab | 34485.53055 | FALSE | TRUE | FALSE | TRUE |
| HLA-A*68:02 | EIVDTVSAL | orf1ab | 17716.53543 | FALSE | TRUE | FALSE | FALSE |
| HLA-B*18:01 | HEFCSQHTM | orf1ab | 9316.770186 | TRUE | TRUE | TRUE | TRUE |
| HLA-B*18:01 | SEYDYVIF | orf1ab | 22818.79195 | TRUE | FALSE | FALSE | FALSE |
| HLA-A*23:01 | VYLPYPDPSRI | orf1ab | 60745.19231 | FALSE | FALSE | FALSE | TRUE |
| HLA-A*23:01 | YLPYPDPSRIL | orf1ab | 31635.67664 | FALSE | FALSE | TRUE | TRUE |
| HLA-A*23:01 | HYVYIGDPAQL | orf1ab | 11972.45717 | FALSE | TRUE | FALSE | TRUE |
| HLA-A*23:01 | LPYPDPSRIL | orf1ab | 20038.18828 | FALSE | FALSE | TRUE | TRUE |
| HLA-A*23:01 | YVYIGDPAQL | orf1ab | 36460.98929 | FALSE | TRUE | FALSE | TRUE |
| HLA-A*23:01 | AYANSVFNI | orf1ab | 10917.03057 | TRUE | FALSE | FALSE | FALSE |
| HLA-A*23:01 | PYPDPSRIL | orf1ab | 20031.26777 | FALSE | FALSE | TRUE | TRUE |
| HLA-A*23:01 | LYLGGMSYY | orf1ab | 11439.33685 | FALSE | TRUE | FALSE | TRUE |
| HLA-A*23:01 | VYIGDPAQL | orf1ab | 212679.4258 | FALSE | TRUE | FALSE | TRUE |
| HLA-A*23:01 | SWNADLYKL | orf1ab | 12169.60352 | FALSE | FALSE | FALSE | TRUE |
| HLA-A*24:02 | VYLPYPDPSRI | orf1ab | 69475.13122 | FALSE | FALSE | FALSE | TRUE |
| HLA-A*24:02 | YLPYPDPSRIL | orf1ab | 44610.20752 | FALSE | FALSE | TRUE | TRUE |
| HLA-A*24:02 | HYVYIGDPAQL | orf1ab | 12054.80948 | FALSE | TRUE | FALSE | TRUE |
| HLA-A*24:02 | LPYPDPSRIL | orf1ab | 28025.09005 | FALSE | FALSE | TRUE | TRUE |
| HLA-A*24:02 | YVYIGDPAQL | orf1ab | 31962.44971 | FALSE | TRUE | FALSE | TRUE |
| HLA-A*24:02 | AYANSVFNI | orf1ab | 14970.05988 | TRUE | FALSE | FALSE | FALSE |
| HLA-A*24:02 | PYPDPSRIL | orf1ab | 25653.44012 | FALSE | FALSE | TRUE | TRUE |

| | | | | | | | |
|---|---|---|---|---|---|---|---|
| HLA-A*24:02 | LYLGGMSYY | orf1ab | 9965.53422 | FALSE | TRUE | FALSE | TRUE |
| HLA-A*24:02 | VYIGDPAQL | orf1ab | 161636.3636 | FALSE | TRUE | FALSE | TRUE |
| HLA-A*24:02 | SWNADLYKL | orf1ab | 14088.39779 | FALSE | FALSE | FALSE | TRUE |
| HLA-A*29:02 | LYLGGMSYY | orf1ab | 165901.6393 | FALSE | TRUE | FALSE | TRUE |
| HLA-A*29:02 | YLGGMSYY | orf1ab | 8398.11543 | FALSE | TRUE | FALSE | TRUE |
| HLA-A*01:01 | QTVDSSQGSEY | orf1ab | 14133.54531 | FALSE | FALSE | TRUE | FALSE |
| HLA-A*01:01 | IVDTVSALVY | orf1ab | 57272.72727 | FALSE | TRUE | FALSE | FALSE |
| HLA-A*01:01 | TVDSSQGSEY | orf1ab | 105494.5055 | FALSE | FALSE | TRUE | FALSE |
| HLA-A*01:01 | DSSQGSEYDY | orf1ab | 12092.71078 | FALSE | FALSE | TRUE | FALSE |
| HLA-A*01:01 | FVDGVPFVV | orf1ab | 11695.747 | FALSE | TRUE | FALSE | TRUE |
| HLA-A*03:01 | ALGGSVAIK | orf1ab | 7374.631268 | FALSE | TRUE | FALSE | FALSE |
| HLA-A*11:01 | YVYLPYPDPSR | orf1ab | 7939.804013 | FALSE | FALSE | FALSE | TRUE |
| HLA-A*30:02 | TVDSSQGSEY | orf1ab | 7399.79445 | FALSE | FALSE | TRUE | FALSE |
| HLA-A*30:02 | LYLGGMSYY | orf1ab | 43841.15523 | FALSE | TRUE | FALSE | TRUE |
| HLA-A*30:02 | SSQGSEYDY | orf1ab | 7801.526718 | FALSE | FALSE | TRUE | FALSE |
| HLA-A*26:01 | EIVDTVSALVY | orf1ab | 119344.2623 | FALSE | TRUE | FALSE | FALSE |
| HLA-A*26:01 | QTVDSSQGSEY | orf1ab | 11196.47355 | FALSE | FALSE | TRUE | FALSE |
| HLA-A*26:01 | EIVDTVSAL | orf1ab | 41925.46584 | FALSE | TRUE | FALSE | FALSE |
| HLA-A*31:01 | YVYLPYPDPSR | orf1ab | 9489.682097 | FALSE | FALSE | FALSE | TRUE |
| HLA-A*31:01 | VYLPYPDPSR | orf1ab | 39843.17825 | FALSE | FALSE | FALSE | TRUE |
| HLA-A*31:01 | WVLNNDYYR | orf1ab | 12262.29508 | FALSE | TRUE | FALSE | TRUE |
| HLA-A*31:01 | YLPYPDPSR | orf1ab | 9822.660577 | TRUE | FALSE | TRUE | TRUE |
| HLA-A*68:01 | YVYLPYPDPSR | orf1ab | 33919.76078 | FALSE | FALSE | FALSE | TRUE |
| HLA-A*68:01 | VYLPYPDPSR | orf1ab | 7715.355805 | FALSE | FALSE | FALSE | TRUE |
| HLA-A*68:01 | WVLNNDYYR | orf1ab | 49500 | FALSE | TRUE | FALSE | TRUE |
| HLA-A*68:01 | YLPYPDPSR | orf1ab | 34376.80667 | TRUE | FALSE | TRUE | TRUE |
| HLA-A*33:01 | YVYLPYPDPSR | orf1ab | 16226.97741 | FALSE | FALSE | FALSE | TRUE |
| HLA-A*33:01 | VYLPYPDPSR | orf1ab | 42620.68966 | FALSE | FALSE | FALSE | TRUE |
| HLA-A*33:01 | WVLNNDYYR | orf1ab | 26671.94929 | FALSE | TRUE | FALSE | TRUE |
| HLA-A*33:01 | YLPYPDPSR | orf1ab | 18690.02496 | TRUE | FALSE | TRUE | TRUE |
| HLA-B*51:01 | VYLPYPDPSRI | orf1ab | 89494.52672 | FALSE | FALSE | FALSE | TRUE |
| HLA-B*51:01 | YLPYPDPSRIL | orf1ab | 25341.29829 | FALSE | FALSE | TRUE | TRUE |
| HLA-B*51:01 | LPYPDPSRILG | orf1ab | 16018.99441 | FALSE | FALSE | FALSE | TRUE |
| HLA-B*51:01 | YLPYPDPSRI | orf1ab | 1009756.098 | TRUE | FALSE | TRUE | TRUE |
| HLA-B*51:01 | LPYPDPSRIL | orf1ab | 248765.1599 | FALSE | FALSE | TRUE | TRUE |
| HLA-B*51:01 | LPYPDPSRI | orf1ab | 5605000 | TRUE | FALSE | TRUE | TRUE |
| HLA-B*51:01 | PYPDPSRI | orf1ab | 29565.753 | TRUE | FALSE | TRUE | TRUE |
| HLA-B*51:01 | YPDPSRIL | orf1ab | 16722.36881 | FALSE | FALSE | TRUE | TRUE |
| HLA-B*35:01 | YLPYPDPSRI | orf1ab | 26136.36364 | TRUE | FALSE | TRUE | TRUE |
| HLA-B*35:01 | LPYPDPSRIL | orf1ab | 39076.89643 | FALSE | FALSE | TRUE | TRUE |
| HLA-B*35:01 | YPDPSRILGA | orf1ab | 8848.641656 | FALSE | FALSE | FALSE | TRUE |
| HLA-B*35:01 | LPYPDPSRI | orf1ab | 59041.43258 | TRUE | FALSE | TRUE | TRUE |
| HLA-B*35:01 | YPDPSRILG | orf1ab | 16681.82717 | FALSE | FALSE | FALSE | TRUE |
| HLA-B*35:01 | YPDPSRIL | orf1ab | 10357.72653 | FALSE | FALSE | TRUE | TRUE |
| HLA-B*35:01 | YAFEHIVY | orf1ab | 17500 | FALSE | TRUE | FALSE | FALSE |
| HLA-B*53:01 | VYLPYPDPSRI | orf1ab | 17794.63543 | FALSE | FALSE | FALSE | TRUE |
| HLA-B*53:01 | YLPYPDPSRIL | orf1ab | 8079.843564 | FALSE | FALSE | TRUE | TRUE |
| HLA-B*53:01 | YLPYPDPSRI | orf1ab | 178064.5161 | TRUE | FALSE | TRUE | TRUE |
| HLA-B*53:01 | LPYPDPSRIL | orf1ab | 85239.89422 | FALSE | FALSE | TRUE | TRUE |

| | | | | | | | |
|---|---|---|---|---|---|---|---|
| HLA-B*53:01 | LPYPDPSRI | orf1ab | 354000 | TRUE | FALSE | TRUE | TRUE |
| HLA-B*53:01 | YPDPSRILG | orf1ab | 12855.83664 | FALSE | FALSE | FALSE | TRUE |
| HLA-B*53:01 | YPDPSRIL | orf1ab | 11741.45739 | FALSE | FALSE | TRUE | TRUE |
| HLA-B*07:02 | VYLPYPDPSRI | orf1ab | 11690.70951 | FALSE | FALSE | FALSE | TRUE |
| HLA-B*07:02 | YLPYPDPSRIL | orf1ab | 14004.13751 | FALSE | FALSE | TRUE | TRUE |
| HLA-B*07:02 | LPYPDPSRILG | orf1ab | 11055.67551 | FALSE | FALSE | FALSE | TRUE |
| HLA-B*07:02 | YLPYPDPSRI | orf1ab | 30799.75201 | TRUE | FALSE | TRUE | TRUE |
| HLA-B*07:02 | LPYPDPSRIL | orf1ab | 184037.5204 | FALSE | FALSE | TRUE | TRUE |
| HLA-B*07:02 | YPDPSRILGA | orf1ab | 9915.096293 | FALSE | FALSE | FALSE | TRUE |
| HLA-B*07:02 | LPYPDPSRI | orf1ab | 60550.95427 | TRUE | FALSE | TRUE | TRUE |
| HLA-B*07:02 | YPDPSRILG | orf1ab | 7536.853391 | FALSE | FALSE | FALSE | TRUE |
| HLA-B*07:02 | YPDPSRIL | orf1ab | 31759.46548 | FALSE | FALSE | TRUE | TRUE |
| HLA-B*54:01 | VYLPYPDPSRI | orf1ab | 16697.90353 | FALSE | FALSE | FALSE | TRUE |
| HLA-B*54:01 | YLPYPDPSRIL | orf1ab | 7030.228036 | FALSE | FALSE | TRUE | TRUE |
| HLA-B*54:01 | LPYPDPSRILG | orf1ab | 9040.292578 | FALSE | FALSE | FALSE | TRUE |
| HLA-B*54:01 | YLPYPDPSRI | orf1ab | 164503.3113 | TRUE | FALSE | TRUE | TRUE |
| HLA-B*54:01 | LPYPDPSRIL | orf1ab | 71765.26718 | FALSE | FALSE | TRUE | TRUE |
| HLA-B*54:01 | YPDPSRILGA | orf1ab | 284154.3027 | FALSE | FALSE | FALSE | TRUE |
| HLA-B*54:01 | FVDGVPFVV | orf1ab | 7903.463522 | FALSE | TRUE | FALSE | TRUE |
| HLA-B*54:01 | LPYPDPSRI | orf1ab | 656835.9375 | TRUE | FALSE | TRUE | TRUE |
| HLA-B*54:01 | YPDPSRILG | orf1ab | 34845.10533 | FALSE | FALSE | FALSE | TRUE |
| HLA-B*54:01 | YPDPSRIL | orf1ab | 8642.424242 | FALSE | FALSE | TRUE | TRUE |
| HLA-B*08:01 | LPYPDPSRIL | orf1ab | 9521.86023 | FALSE | FALSE | TRUE | TRUE |
| HLA-B*08:01 | YLRKHFSMM | orf1ab | 47421.38365 | FALSE | FALSE | TRUE | FALSE |
| HLA-B*08:01 | LPYPDPSRI | orf1ab | 9395.954403 | TRUE | FALSE | TRUE | TRUE |
| HLA-B*08:01 | SLYVNKHAF | orf1ab | 9338.374291 | FALSE | TRUE | TRUE | TRUE |
| HLA-B*08:01 | YLRKHFSM | orf1ab | 53164.55696 | FALSE | FALSE | TRUE | FALSE |
| HLA-B*08:01 | YPDPSRIL | orf1ab | 8265.468773 | FALSE | FALSE | TRUE | TRUE |
| HLA-C*06:02 | YLPYPDPSRI | orf1ab | 15017.22991 | TRUE | FALSE | TRUE | TRUE |
| HLA-C*06:02 | LPYPDPSRI | orf1ab | 9673.243974 | TRUE | FALSE | TRUE | TRUE |
| HLA-C*06:02 | LYLGGMSYY | orf1ab | 9494.917905 | FALSE | TRUE | FALSE | TRUE |
| HLA-C*06:02 | VYIGDPAQL | orf1ab | 13073.52941 | FALSE | TRUE | FALSE | TRUE |
| HLA-A*69:01 | YLPYPDPSRI | orf1ab | 14531.41453 | TRUE | FALSE | TRUE | TRUE |
| HLA-A*69:01 | YVYIGDPAQL | orf1ab | 12619.13166 | FALSE | TRUE | FALSE | TRUE |
| HLA-A*69:01 | FVDGVPFVV | orf1ab | 184120.1717 | FALSE | TRUE | FALSE | TRUE |
| HLA-A*69:01 | LPYPDPSRI | orf1ab | 10195.85253 | TRUE | FALSE | TRUE | TRUE |
| HLA-A*69:01 | EIVDTVSAL | orf1ab | 14900.66225 | FALSE | TRUE | FALSE | FALSE |
| HLA-B*15:01 | VLYYQNNVF | orf1ab | 18967.97153 | FALSE | TRUE | FALSE | TRUE |
| HLA-B*15:01 | SLYVNKHAF | orf1ab | 15901.28755 | FALSE | TRUE | TRUE | TRUE |
| HLA-C*14:02 | VYLPYPDPSRI | orf1ab | 7543.48829 | FALSE | FALSE | FALSE | TRUE |
| HLA-C*14:02 | YLPYPDPSRIL | orf1ab | 14219.44134 | FALSE | FALSE | TRUE | TRUE |
| HLA-C*14:02 | YLPYPDPSRI | orf1ab | 7433.119876 | TRUE | FALSE | TRUE | TRUE |
| HLA-C*14:02 | LPYPDPSRIL | orf1ab | 13025.63214 | FALSE | FALSE | TRUE | TRUE |
| HLA-C*14:02 | YVYIGDPAQL | orf1ab | 23706.89655 | FALSE | TRUE | FALSE | TRUE |
| HLA-C*14:02 | IFVDGVPFV | orf1ab | 13494.64224 | FALSE | TRUE | FALSE | TRUE |
| HLA-C*14:02 | LYYQNNVFM | orf1ab | 23333.33333 | FALSE | TRUE | TRUE | TRUE |
| HLA-C*14:02 | LPYPDPSRI | orf1ab | 7171.493155 | TRUE | FALSE | TRUE | TRUE |
| HLA-C*14:02 | LYLGGMSYY | orf1ab | 47400.46838 | FALSE | TRUE | FALSE | TRUE |
| HLA-C*14:02 | VYIGDPAQL | orf1ab | 180691.0569 | FALSE | TRUE | FALSE | TRUE |

| | | | | | | | |
|---|---|---|---|---|---|---|---|
| HLA-C*14:02 | SLYVNKHAF | orf1ab | 7658.914729 | FALSE | TRUE | TRUE | TRUE |
| HLA-C*14:02 | SWNADLYKL | orf1ab | 7661.197134 | FALSE | FALSE | FALSE | TRUE |
| HLA-B*15:42 | YLPYPDPSRI | orf1ab | 12438.03515 | TRUE | FALSE | TRUE | TRUE |
| HLA-B*15:42 | LPYPDPSRIL | orf1ab | 8421.858087 | FALSE | FALSE | TRUE | TRUE |
| HLA-B*15:42 | LPYPDPSRI | orf1ab | 37500 | TRUE | FALSE | TRUE | TRUE |
| HLA-B*15:42 | LVYDNKLKA | orf1ab | 7701.746107 | FALSE | FALSE | FALSE | TRUE |
| HLA-B*15:42 | GVAPGTAVL | orf1ab | 7268.207805 | TRUE | FALSE | FALSE | FALSE |
| HLA-B*45:06 | YLPYPDPSRI | orf1ab | 9013.062409 | TRUE | FALSE | TRUE | TRUE |
| HLA-B*45:06 | YPDPSRILGA | orf1ab | 13625.49801 | FALSE | FALSE | FALSE | TRUE |
| HLA-B*45:06 | LPYPDPSRI | orf1ab | 26703.192 | TRUE | FALSE | TRUE | TRUE |
| HLA-B*45:06 | YPDPSRILG | orf1ab | 7974.288685 | FALSE | FALSE | FALSE | TRUE |
| HLA-C*04:01 | YLPYPDPSRIL | orf1ab | 34166.66667 | FALSE | FALSE | TRUE | TRUE |
| HLA-C*04:01 | IFVDGVPFVV | orf1ab | 31594.75465 | FALSE | TRUE | FALSE | TRUE |
| HLA-C*04:01 | YLPYPDPSRI | orf1ab | 19270.75252 | TRUE | FALSE | TRUE | TRUE |
| HLA-C*04:01 | LPYPDPSRIL | orf1ab | 17837.77374 | FALSE | FALSE | TRUE | TRUE |
| HLA-C*04:01 | YVYIGDPAQL | orf1ab | 9785.137539 | FALSE | TRUE | FALSE | TRUE |
| HLA-C*04:01 | IFVDGVPFV | orf1ab | 29464.15094 | FALSE | TRUE | FALSE | TRUE |
| HLA-C*04:01 | FVDGVPFVV | orf1ab | 56078.43137 | FALSE | TRUE | FALSE | TRUE |
| HLA-C*04:01 | LPYPDPSRI | orf1ab | 11830.71836 | TRUE | FALSE | TRUE | TRUE |
| HLA-C*04:01 | PYPDPSRIL | orf1ab | 10605.76417 | FALSE | FALSE | TRUE | TRUE |
| HLA-C*04:01 | VYIGDPAQL | orf1ab | 17576.11704 | FALSE | TRUE | FALSE | TRUE |
| HLA-C*04:01 | SWNADLYKL | orf1ab | 14154.56874 | FALSE | FALSE | FALSE | TRUE |
| HLA-C*04:01 | WYDFGDFI | orf1ab | 13264.9388 | FALSE | FALSE | FALSE | TRUE |
| HLA-C*04:01 | YPDPSRIL | orf1ab | 8671.328671 | FALSE | FALSE | TRUE | TRUE |
| HLA-C*15:02 | YLPYPDPSRI | orf1ab | 17093.31131 | TRUE | FALSE | TRUE | TRUE |
| HLA-C*15:02 | YVYIGDPAQL | orf1ab | 12323.33678 | FALSE | TRUE | FALSE | TRUE |
| HLA-C*15:02 | FVDGVPFVV | orf1ab | 175819.6721 | FALSE | TRUE | FALSE | TRUE |
| HLA-C*15:02 | LPYPDPSRI | orf1ab | 10556.21822 | TRUE | FALSE | TRUE | TRUE |
| HLA-C*15:02 | GVAPGTAVL | orf1ab | 23187.94607 | TRUE | FALSE | FALSE | FALSE |
